# How and why plants and human N-glycans are different: Insight from molecular dynamics into the “glycoblocks” architecture of complex carbohydrates

**DOI:** 10.1101/2020.05.22.110528

**Authors:** CA Fogarty, AM Harbison, AR Dugdale, E Fadda

## Abstract

N-glycosylation is one of the most abundant and diverse post-translational modifications of proteins, implicated in protein folding and structural stability, and mediating interactions with receptors and with the environment. All N-glycans share a common core from which linear or branched arms stem from, with functionalization specific to different species and to the cells’ health and disease state. This diversity generates a rich collection of structures, all diversely able to trigger molecular cascades and to activate pathways, which also include adverse immunogenic responses. These events are inherently linked to the N-glycans 3D architecture and dynamics, which remain for the large part unresolved and undetected because of their intrinsic structural disorder. In this work we use molecular dynamics (MD) simulations to provide insight into N-glycans 3D structure by analysing the effects of a set of very specific modifications found in plants and invertebrate N-glycans, which are immunogenic in humans. We also compare these structural motifs and combine them with mammalian N-glycans motifs to devise strategies for the control of the N-glycan 3D structure through sequence. Our results suggest that the N-glycans architecture can be described in terms of the local spatial environment of groups of monosaccharides. We define these “glycoblocks” as self-contained 3D units, uniquely identified by the nature of the residues they comprise, their linkages and structural/dynamic features. This alternative description of glycans 3D architecture can potentially lead to an easier prediction of sequence-to-structure relationships in complex carbohydrates, with important implications in glycoengineering design.

## Introduction

Complex carbohydrates (or glycans) are an essential class of biomolecules, directly implicated in the cell’s interactions with its environment, facilitating communication and infection[1,2]. These processes are often initiated by molecular recognition involving carbohydrate-binding proteins (lectins) or by glycan-glycan interactions[1,3-5], all events that hinge on specific structural and dynamic features of the glycans. This makes the 3D complementarity of the glycans architecture key towards the success of these processes and an essential piece of information for us to have in order to understand glycan recognition. Because of their chemical nature, glycans are intrinsically flexible and highly dynamic at room temperature, thus their characterization through experimental structural biology methods is hardly straightforward even in cryogenic environments[6]. As an additive layer of difficulty, glycosylation is only indirectly dependent on the genome, which often results in a micro- (or macro-)heterogeneity of glycan sequences at specific sites[7]. These complexities are very difficult to resolve, requiring high levels of expertise and multi-layered orthogonal approaches[8-10,7]. Within this framework, the contribution of glycoinformatics tools and databases represents an essential resource to advance glycomics[11-15], while molecular simulations fit in very well as complementary and orthogonal techniques to support and advance structural glycobiology research. Indeed, current high performance computing (HPC) technology allows us to study realistic model systems[16,17] and to reach experimental timescales[18], so that computing can now contribute as one of the leading research methods in structural glycobiology.

One of the most interesting and remarkably challenging areas in glycoscience research that HPC simulations can address is the study of the links between glycans sequence and 3D structure. This direct relationship is a well-recognized and broadly accepted concept in proteins’ structural biology, according to which the amino acid sequence dictates the functional 3D fold and its stability. However, the same notion is not generally invoked when discussing other biopolymers or complex carbohydrates. In the specific case of glycans, the structural complexity, in terms of the diversity of monosaccharides, the linkages’ stereochemistry and the branched scaffolds, makes the already difficult case even more intricate. Nevertheless, the fact that glycoforms follow recurrent sequence patterns, clearly suggests that the glycans 3D structure is also non-random and very likely sequence-determined. We use computer modelling to gain insight into these relationships and to define a framework to understand how subtle modifications to the glycans sequence can alter their 3D structure and conformational dynamics, ultimately regulating recognition[19]. In this work we use molecular dynamics (MD) simulations to analyse the effects of the inclusion of motifs typically found in plants and invertebrates N-glycans and immunogenic in mammals[20-23]. More specifically, we investigate how core α(1-3)-linked fucose (Fuc) and β(1-2)-linked xylose (Xyl) affect the structure and dynamics of plants N-glycoforms[23] and of hybrid constructs with mammalian N-glycoforms[24].

At first glance plants protein N-glycosylation[23] is quite similar to the one of higher species[25], carrying the distinctive trimannose core (Man3), which can be further functionalised with β(1-2) linked GlcNAc residues on the arms. As a trademark feature, shown in **Figure 1**, plants N-glycans can also have a β(1-2)-Xyl linked to the central mannose and core α(1-3)-Fuc, instead of the α(1-6)-Fuc commonly found in mammalian complex N-glycans. Additionally, the arms can be further functionalised with terminal galactose (Gal) in β(1-3) instead of β(1-4)[23], commonly found in vertebrates, which forces the addition of fucose in the α(1-4) position of the GlcNAc and results in the occurrence of Lewis A (LeA) instead of Lewis X (LeX) terminal motifs on the arms[26,23]. In a previous study, we characterized through extensive sampling the structure and dynamics of complex biantennary N-glycans commonly found in the human IgGs Fc region[24]. The results of this study indicated a clear sequence-to-structure relationships, especially in the context of the dynamics of the (1-6) arm. More specifically, we found that the outstretched (open) conformation of the (1-6) arm gets progressively less populated as the functionalization of the arm grows, i.e. from 85% in Man3, to 52% in (F)A2, (F)A2[3]G1, and (F)A2[3]G1S1, where the (F) indicates the presence or absence of α(1-6) core fucosylation, to 24% in all structures with (1-6) arm terminating with Gal-β(1-4)-GlcNAc or Sia-α(2-6)-Gal-β(1-4)-GlcNAc, irrespective of the functionalization of the (1-3) arm[24]. As a practical implication of these results, positional isomers, such as (F)A2[3]G1 and (F)A2[6]G1, have different conformational propensities, the latter with a much lower population of outstretched (1-6) arm and therefore quite different 3D average structures, which ultimately explains their differential recognition in glycan arrays[27]. Additionally, the different conformation of the arms explains the known difficulties in sialylating the (1-6) arm by ST6-Gal1, relatively to the (1-3) arm[28]. Also, the different 3D conformational propensity of the arms in function of sequence can have important implications in terms of the N-glycans biosynthesis and biodegradation[29]. As an additional interesting point, we found that the folding of the (1-6) arm over the chitobiose region is completely independent of core α(1-6) fucosylation[24], with the result that core-fucosylated and non-core fucosylated N-glycans with the same sequence in the (1-6) arm correspond to the same structural ensemble.

**Figure 1.**
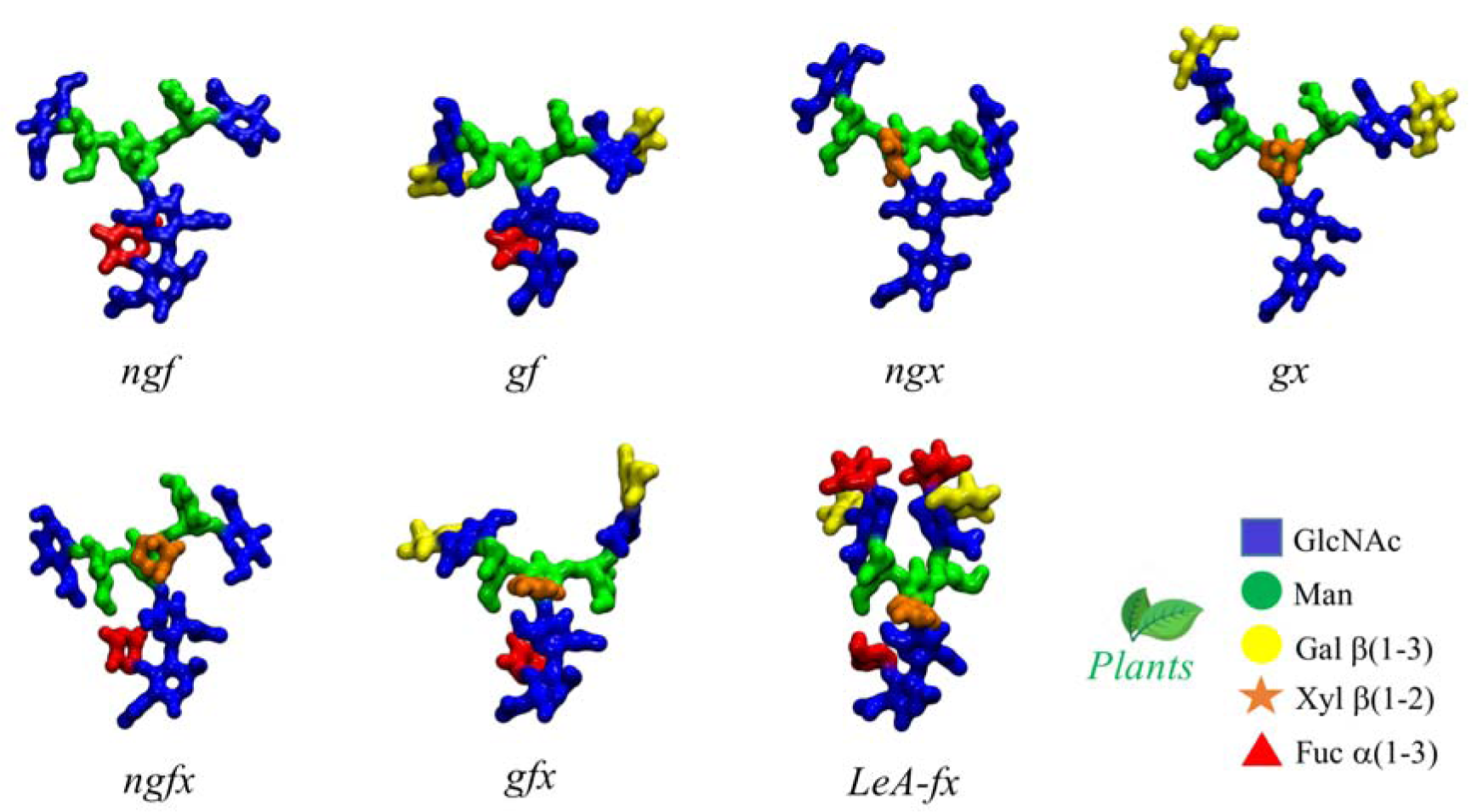
Representative structures of the plant N-glycans studied in this work with corresponding nomenclature. The letters *f, x*, and *g* indicate the presence of Fuc, Xyl and β(1-3) Gal, respectively, and *ng* the absence of β(1-3) Gal. *LeA* stands for Lewis A antigen. The N-glycans structures are shown with the (1-3) and (1-6) arms on the left and on the right, respectively. The monosaccharides colouring follows the SFNG nomenclature. The plants N-glycan characteristic linkages are indicated in the legend. Rendering was done with VMD (http://www.ks.uiuc.edu/Research/vmd/).

In this work we discuss how core α(1-3)-Fuc and β(1-2)-Xyl regulate the conformational propensity of the (1-6) arm to push a predominantly outstretched (open) conformation when the arms are functionalized with terminal β(1-3)-Gal. Within this framework, we explored the possibility of integrating these motifs in the context of mammalian sequences as an exploratory strategy towards the design of N-glycans with the desired 3D structure. For simplicity in the presentation and discussion of the results, we refer to N-glycans as either “plant” or “hybrid” separately. Nevertheless, it is important to underline that some of these motifs, such as β(1-2) xylosylation and difucosylated core are also found in invertebrate N-glycosylation[30]. Finally, we discuss these findings within a framework where the different N-glycoforms can be represented as a combination of spatial self-contained units, named “glycoblocks”, rather than in terms of monosaccharides and linkages. We find that this approach helps our understanding of N-glycans architecture in terms of equilibrium structures and relative populations and also of how specific modifications affect molecular recognition.

## Computational Methods

All starting structures were generated with the GLYCAM Carbohydrate Builder (http://www.glycam.org). For each sequence we selected the complete set of torsion angle values obtained by variation of the 1-6 dihedrals, namely the three *gg, gt* and *tg* conformations for each 1-6 torsion. The topology file for each structure was obtained using *tleap*[31], with parameters from the GLYCAM06-j1[32] for the carbohydrate atoms and with TIP3P for water molecules[33]. All calculations were run with the AMBER18 software package[31] on NVIDIA Tesla V100 16GB PCIe (Volta architecture) GPUs installed on the HPC infrastructure *kay* at the Irish Centre for High-End Computing (ICHEC). Separate production steps of 500 ns each were run for each rotamer (starting system) and convergence was assessed based on conformational and clustering analysis, see Supplementary Material for all relevant Tables. Simulations were extended, if the sampling was not deemed sufficient, i.e. in case standard deviation values measured were significantly larger than 15° for each cluster in each trajectory. All trajectories were processed using *cpptraj*[31] and visually analysed with the Visual Molecular Dynamics (VMD) software package[34]. Backbone Root Mean Square Deviation (RMSD) and torsion angles values were measured using VMD. A density-based clustering method was used to calculate the populations of occupied conformations for each torsion angle in a trajectory and heat maps for each dihedral were generated with a kernel density estimate (KDE) function. Statistical and clustering analysis was done with the R package and data were plotted with RStudio (www.rstudio.com). Further details on the simulation set-up and running protocol are included as Supplementary Material.

## Results

### Core α(1-3) fucose in plant N-glycans

One distinctive feature of plants N-glycans is the occurrence of core fucosylation in α(1-3), rather than α(1-6)-Fuc, normally found in mammalian N-glycans[24,23]. To understand the effects on the 3D structure of this modification, we have considered two biantennary systems, one terminating with β(1-2)-GlcNAc on both arms (*ngf*) and the other with terminal β(1-3)-Gal on both arms (*gf*), shown in **Figure 1**. In both glycoforms core α(1-3)-Fuc occupies a stable position, with one single conformer populated (100%), see **Tables S.1 and S.2**. This conformation is supported by a stacking interaction between the core α(1-3)-Fuc and β(1-4) GlcNAc of the chitobiose in a “closed” conformation, which resembles the stable conformation of LeX[35]. This spatial arrangement imposes a 20° rotation of the GlcNAc-β(1-4)-GlcNAc linkage, see **Tables S.1** and **S.2**, relative to the α(1-6) core fucosylated or non-fucosylated chitobiose[24], where the average psi value is −127.8° (14.8)[24], but doesn’t affect the structure of the linkage to the central mannose. As shown by the low standard deviation values and by the lack of multiple minima (clusters), the N-glycan core remains relatively rigid throughout the trajectories. The slight torsion of the GlcNAc-β(1-4)-GlcNAc linkage imposed by the α(1-3)-Fuc has a dramatic effect on the conformational dynamics of the (1-6) arm, which is found predominantly in an outstretched (66%, cluster 1) conformation, rather than folded over (34%, clusters 1 and 2), see **Table S.1**. The addition of a terminal β(1-3)-Gal in the *gf* N-glycan pushes the equilibrium towards an outstretched (1-6) arm even further, with the open conformation populated at 72%, see **Table S.2**. Interestingly, in the case of α(1-6) core fucosylated N-glycans, and with double fucosylation as discussed later on, the equilibrium of the (1-6) arm was the exact opposite, with a predominance of the folded conformation, especially in the presence of terminal β(1-4) Gal[24]. To note, the folded (1-6) arm conformation can be either a ‘front fold’, see **Figure 2 panel a**, where the torsion around the α(1-6) linkage brings the arm towards the reader, or a ‘back fold’ where the (1-6) arm interacts with the α(1-3)-Fuc, away from the reader. As shown in **Tables S.1** and **S.2**, the equilibrium of the (1-3) arm is not affected by core α(1-3)-Fuc.

**Figure 2.**
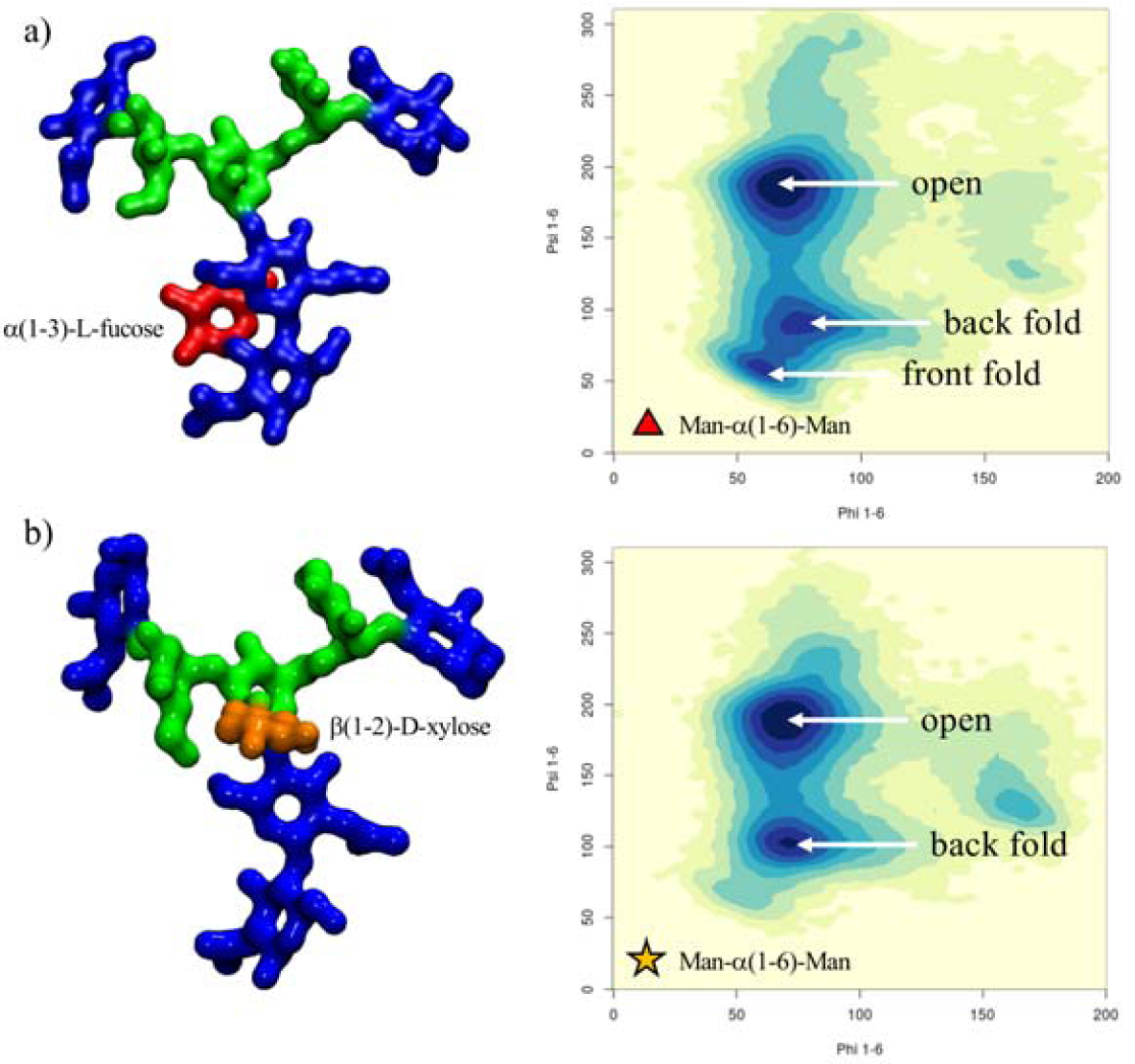
A representative structure of the non-galactosylated N-glycan with α(1-3)-linked core fucose (*ngf*) is shown in panel a), with on the right-hand side the corresponding heat map showing the conformations accessible to the (1-6) arm in terms of the phi/psi torsion angles. A representative structure of the non-galactosylated N-glycan with β(1-2)-linked xylose (*ngx*) is shown in panel b), with on the right-hand side a heat map showing the conformations accessible to the (1-6) arm in terms of the phi/psi torsion angles. The N-glycans structures are shown with the (1-3) and (1-6) arms on the left and on the right, respectively. The monosaccharides colouring follows the SFNG nomenclature. The structure rendering was done with VMD and the graphical statistical analysis with *RStudio* (www.rstudio.com).

#### β(1-2) xylose in plant N-glycans

Because the β(1-2)-Xyl sits in front of the two arms, it greatly affects their dynamics. Because of steric hindrance, the (1-3) arm is much more rigid relative to non-xylosylated species, see **Table S.3**, losing its “two conformer” dynamics characteristic of the biantennary mammalian N-glycans[24], also retained in the plant N-glycans with only α(1-3)-Fuc discussed above, see also **Tables S.1** and **S.2**. In regards to the (1-6) arm, as shown in **Figure 2 panel b**, tshe presence of β(1-2)-Xyl has a very similar effect as the α(1-3)-Fuc, pushing the equilibrium towards an open conformation. To note, in the presence β(1-2)-Xyl, the (1-6) arm cannot fold over the chitobiose core in a ‘front fold’ either, because of steric hindrance. Also, similarly to the α(1-3) fucosylated glycans, the stability of the open structure is slightly increased when the arm is further functionalized with terminal β(1-3)-Gal, see **Table S.4**. As an additional interesting feature, through the cumulative 3 μs MD sampling, the xylose ring repeatedly inverts its conformation from the all equatorial ^4^C_1_ chair, to the ^1^C_4_ chair, where all hydroxyl groups are axial, see **Figure 3**. This transition may be energetically facilitated by the hydrogen bonding interaction xylose is able to form when in a ^1^C_4_ chair with the α(1-6)-Man, which may compensate for the steric compression, making the ^1^C_4_ chair the highest populated conformer at 76% within an N-glycan scaffold. Both experimental and *ab-initio* theoretical studies[36-38] have shown that ^1^C_4_ chair is energetically accessible in isolated β-D-Xyl at room temperature in different dielectric conditions.

**Figure 3.**
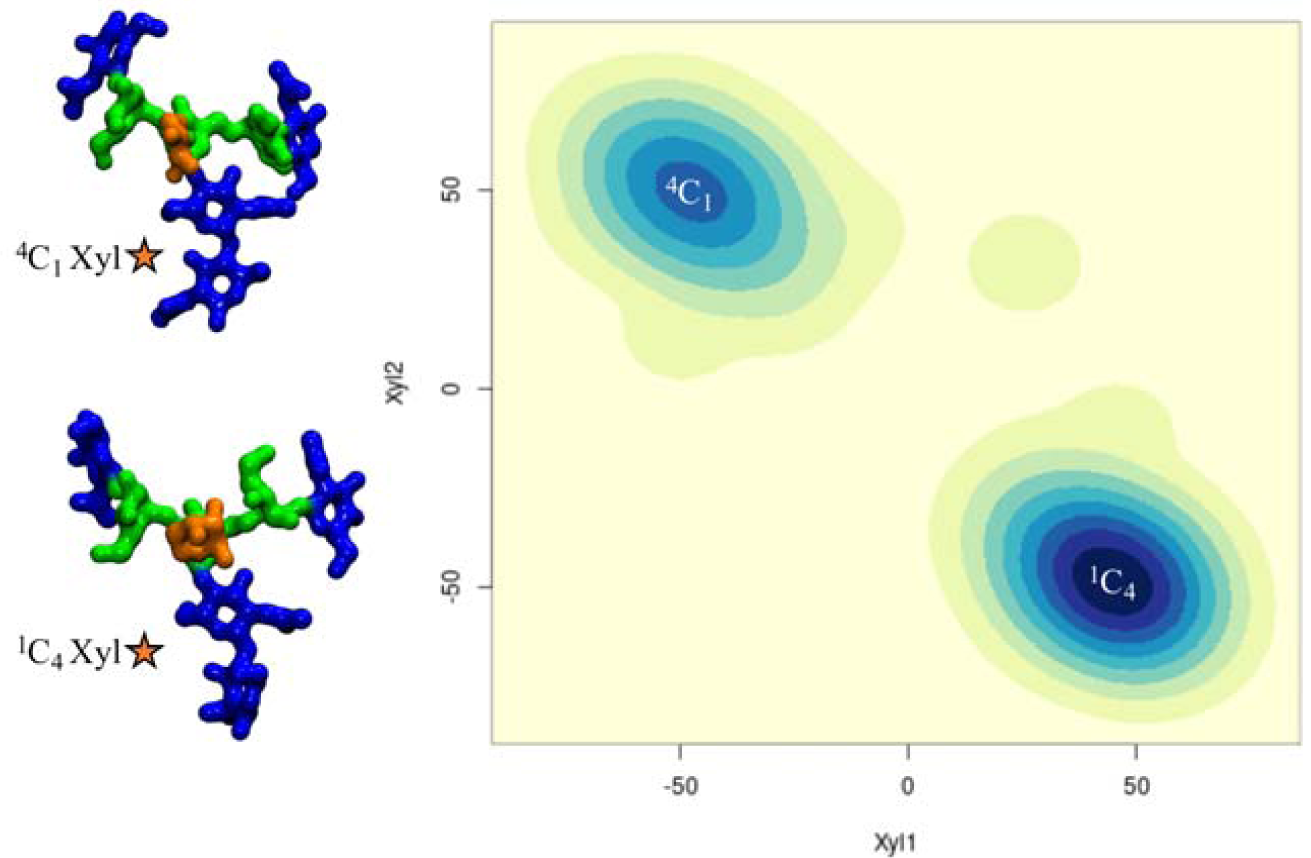
β-D-xylose ring pucker analysis over 3 μs of cumulative MD sampling of the *ngx* N-glycan. The two snapshots on the right-hand side are representative *ngx* conformations corresponding to the two different ring puckers. The Xyl1 and Xyl2 axis labels refer to the torsion angles C1C2C3C4 and C2C3C4C5, respectively. The N-glycans structures are shown with the (1-3) and (1-6) arms on the left and on the right, respectively. The monosaccharides colouring follows the SFNG nomenclature. The structure rendering was done with VMD and the graphical statistical analysis with *RStudio* (www.rstudio.com).

#### Core α(1-3) fucose and β(1-2) xylose in plant N-glycans

The presence of both α(1-3)-Fuc and β(1-2)-Xyl brings in the characteristic features highlighted earlier in the analysis of the structures with either α(1-3)-Fuc or β(1-2)-Xyl. Indeed, we see here again the 20° rotation of the chitobiose GlcNAc-β(1-4)-GlcNAc *psi* angle caused by the stacking of the α(1-3)-Fuc to the chitobiose β(1-4)-GlcNAc and the conformational restraints imposed by the β(1-2)-Xyl on the (1-3) arm, see **Table S.5**. We also observed that both α(1-3)-Fuc and β(1-2)-Xyl push the (1-6) arm equilibrium towards an open conformation, which is also the case when both are present in the *ngfx* N-glycan and to an even higher degree, i.e. 87%, in the *gfx* N-glycan, when both arms are functionalized with terminal β(1-3)-Gal, see **Table S.6**. One feature specific to the *ngfx* N-glycan is the higher flexibility of the core Man-β(1-4)-GlcNAc linkage, which allows for the rotation of the trimannose group relative to the chitobiose core. This conformation was accessible, but only populated around 2% when either β(1-2)-Xyl or α(1-3)-Fuc are present, see **Tables S.1** to **S.4**. When both fucose and xylose are present, the population of the rotated trimannose reaches above 20%, see **Table S.5**, which can be considered a synergistic effect as this conformation is stabilized by a hydrogen bonding network involving the core fucose, the GlcNAc on the (1-6) arm and the xylose, as shown in **Figure S.1**. Such folding event has been observed as a stable conformation in two independent simulations. To note, the functionalization of the arms to include terminal β(1-3)-Gal reduces the occurrence of this event down to around 5%, see **Table S.6**.

#### Terminal LeA and LeX motifs in plant N-glycans

To understand how an increased complexity on the arms would affect the dynamics of the α(1-3) fucosylated and β(1-2) xylosylated N-glycans, we considered the functionalization with terminal LeA antigens present in plants N-glycans[26] and with LeX for comparison. As expected[35] the LeA and LeX structures are quite rigid, see **Tables S.7** and **S.15**, and remain in what is known as the “closed” conformation throughout the 1.5 μs cumulative sampling time for each system. One interesting point is that the branching introduced by functionalizing the terminal GlcNAc residues with α(1-4)-Fuc and β(1-3)-Gal, i.e. LeA, promotes the interaction between the two arms, which is not observed when the arms are linear, neither here for plants N-glycans, nor for mammalian IgG-type complex biantennary N-glycans[24]. The interaction between the arms is promoted by the ability to form complex hydrogen bonding networks, which in this specific case, may also involve the central xylose. As outcomes of the complex interaction the branched arms can establish, the equilibrium of the (1-6) arm is restrained in conformations previously not significantly populated, see **Figure 4** and **Table S.7**, and the GlcNAc-β(1-2)-Man linkage in both arms is remarkably flexible, which is also not observed when the arms are not branched. Although not natural in plants, to check the corresponding symmetry, we built a core α(1-3)-Fuc and β(1-2)-Xyl N-glycan with terminal LeX on both arms, a feature actually found in schistosome N-glycosylation[30]. Remarkably, as shown in **Figure 4** and **Table S.15**, within this framework the dynamics of the (1-6) arm is completely different. Contrary to the N-glycan with terminal LeA groups, the two arms with LeX are not interacting and the (1-6) arm is predominantly (90%) in an extended (open) conformation, while the closed conformation, which accounts for the remaining 10% is achieved through a rotation around the core Man-β(1-4)-GlcNAc. The lack of interaction between the arms is due to the inability to establish the same stable hydrogen bonding network due to the non-complementary position of the deoxy-C6 of the fucose in LeX relative to LeA.

**Figure 4.**
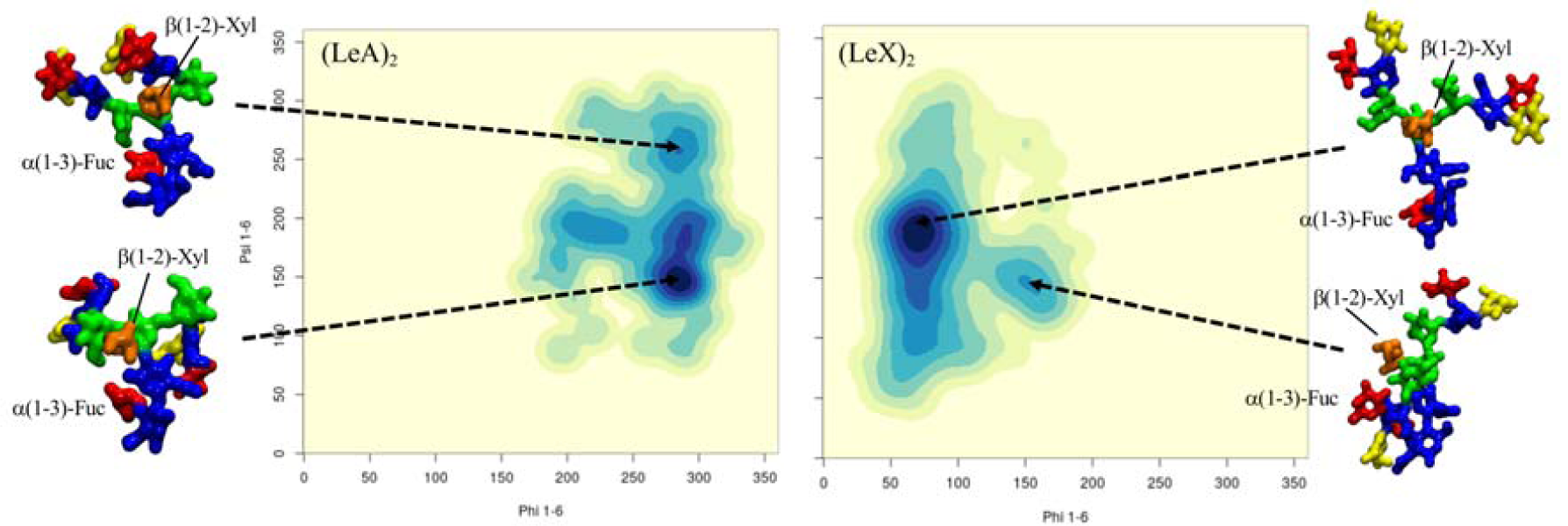
Comparison of the different conformational equilibria of the (1-6) arm in a core α(1-3)-Fuc β(1-2)-Xyl A2 N-glycan with terminal LeA and LeX groups on the left- and right-hand side, respectively. Representative structures from 1.5 μs MD sampling of each system are shown to illustrate the conformations corresponding to the different minima. The N-glycans structures are shown with the (1-3) and (1-6) arms on the left and on the right, respectively. The monosaccharides colouring follows the SFNG nomenclature. The structure rendering was done with VMD and the graphical statistical analysis with *RStudio* (www.rstudio.com).

#### Hybrid N-glycans

To understand how characteristic plant N-glycan motifs can affect the structure of mammalian N-glycoforms, we have designed and analysed the dynamics of a set of hybrid systems. In particular, we were interested in the effect of the addition of β(1-2)-Xyl and α(1-3)-Fuc to (F)A2G2 N-glycans scaffolds in terms of potential alteration of the (1-6) arm dynamics.

#### β(1-2)-xylosylated mammalian N-glycans

Unlike the case of plants N-glycans, the presence of β(1-2)-Xyl hinders but does not completely prevent the (1-6) arm from folding over when the terminal galactose is β(1-4)-linked, as folding over the chitobiose can be stabilized by stacking, see **Figure 5** and **Table S.8**. The folded conformation with a median *psi* value of 103.5° (± 11.3) is 20° from the average value of 82.9° calculated for the non-xylosylated (mammalian) counterpart[24], so slightly distorted, and its population reduced from 74% to 57%. Nevertheless, the closed conformation is still the predominant form, even with β(1-2)-Xyl. The presence of α(1-6)-linked core fucose to create a β(1-2)-xylosylated FA2G2, which is actually a type of N-glycosylation found in schistosoma[30], brings in yet another change. As shown in **Figure 5** and **Table S.9**, α(1-6)-Fuc and β(1-2)-Xyl are in an optimal conformation to support the closed (folded) (1-6) arm, by stacking of the terminal galactose by fucose and hydrogen bonding by xylose. Within this context the closed (1-6) arm is the highest populated conformer at 70.0% over 4.5 μs of cumulative sampling of this system. To note that the conformation of the α(1-6)-linked core fucose is the same as the one seen in mammalian N-glycans[24], which on its own we have seen is not enough to affect the (1-6) arm equilibrium, see **Table S.9**. The interaction of the α(1-6)-Fuc with the terminal β(1-4)-Gal is essential to promote the closed conformation of the (1-6) arm as demonstrated by the results obtained for the xylosylated FA2 systems, which recovers a conformational propensity similar to the non-fucosylated, xylosylated A2G2, see **Figure 5** and **Tables S.8** and **S.10**.

**Figure 5.**
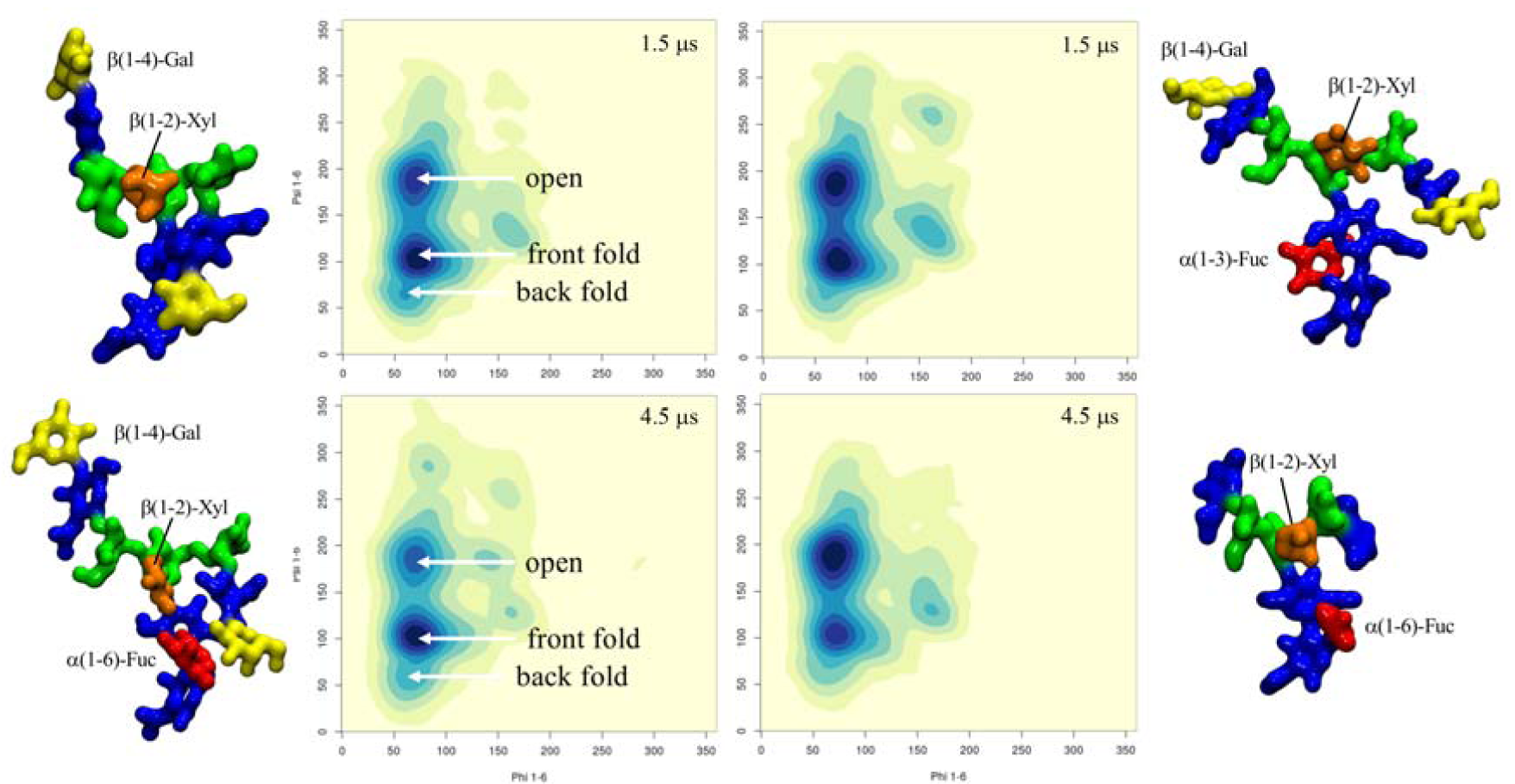
Conformational analysis of the (1-6) arm in four hybrid N-glycoforms, β(1-2)-xylosylated A2G2 (top-left), β(1-2)-xylosylated FA2G2 (bottom-left), β(1-2)-xylosylated α(1-3)-core fucosylated A2G2 (top-right) and β(1-2)-xylosylated FA2 (bottom-right). The predominant conformations are indicated in the top- and bottom-left heat maps for simplicity. The simulation time relative to each system is indicated in the top right corner of each heat map. The N-glycans structures are shown with the (1-3) and (1-6) arms on the left and on the right, respectively. The monosaccharides colouring follows the SFNG nomenclature. The structure rendering was done with VMD and the graphical statistical analysis with *Rstudio* (www.rstudio.com).

#### α(1-3)-fucosylated mammalian N-glycans

Because of its orientation tucked “behind” the chitobiose core defined in the context of plants N-glycans earlier, the effect of core α(1-3)-Fuc on the (1-6) arm equilibrium within an A2G2-xylosylated scaffold is not as significant as α(1-6)-Fuc. As shown in **Figure 5** and **Table S.11**, this lack of direct effect is demonstrated by the recovery of the same equilibrium as the non-fucosylated A2G2-xylosylated system. The dynamics of the chitobiose core is very similar to the one determined for the corresponding plant N-glycan. To analyse the effect of core α(1-3) fucosylation without β(1-2)-Xyl, we have looked at two A2G2 hybrid systems, one with only α(1-3)-linked fucose and one with both core α(1-3)-and α(1-6)-linked fucose, a characteristic “double-fucose” glycosylation found in worm and fly cells[30]. As shown in **Table S.12** unlike in plants N-glycans, the α(1-3)-Fuc alone does not affect the A2G2 (1-6) arm equilibrium[24], as the folding of the (1-6) arm with terminal β(1-4)-Gal is not obstructed by the rotation of the chitobiose core imposed by the α(1-3)-Fuc position. When both α(1-3)- and α(1-6)-linked fucoses are present the (1-6) arm with terminal β(1-4)-Gal is predominantly folded (closed) at 85%, see **Figure 6** and **Table S.13**, which is higher than in the absence of α(1-3)-Fuc[24]. Indeed, the latter can actively contribute in stabilizing the interaction with the terminal β(1-4)-Gal of the folded (1-6) arm. We also observed interesting events, one representing 10% of 2 μs as indicated by the values of the GlcNAc-β(1-4)-GlcNAc torsion, where the GlcNAc is stacked in between the two fucose residues and another one, contributing to 18% of the simulation time, 14% when the system is also xylosylated, in which the GlcNAc ring transitions from ^4^C_1_ to ^1^C_4_ allowing the two fucose to stack, see **Tables S.13** and **S.14** and **Figure S.2**.

**Figure 6.**
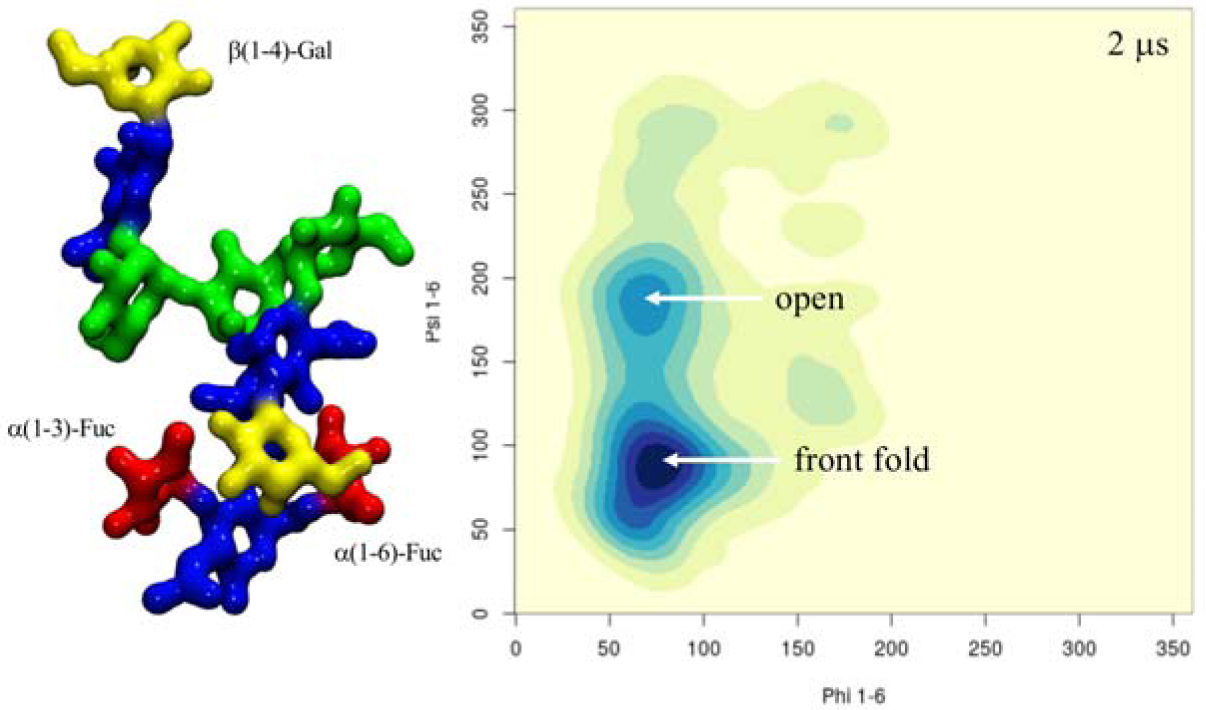
Conformational equilibrium of the (1-6) arm in terms of phi/psi torsion angle values for the α(1-3)-fucosylated FA2G2 N-glycoform. The structure with the folded (1-6) arm where the terminal β(1-4)-Gal interacts with both fucose residues is shown on the left-hand side. The N-glycans structures are shown with the (1-3) and (1-6) arms on the left and on the right, respectively. The monosaccharides colouring follows the SFNG nomenclature. The structure rendering was done with VMD and the graphical statistical analysis with *RStudio* (www.rstudio.com).

## Discussion

Differences and similarities in N-glycans sequences are highly cell-specific as well as important indicators of health and disease states[1,39]. Exogenous N-glycans motifs can be quite subtle, yet trigger profound differences in terms of molecular recognition[19,27] and dangerous immunogenic responses[20-22]. In this work we have analysed the effects on the N-glycans structure and dynamics of two motifs in particular, namely β(1-2)-Xyl and core α(1-3)-Fuc, common in plants[23] and invertebrates[30], but completely absent in mammalian N-glycans. Within the context of plant-type N-glycans, which have a terminal β(1-3)-Gal, rather than β(1-4)-Gal, both β(1-2)-Xyl and α(1-3)-Fuc contribute independently in promoting an outstretched (open) conformation of the (1-6) arm because of steric hindrance of the xylose and of the rotation forced upon the chitobiose core by the α(1-3)-Fuc. The latter is not an obstruction for the folding of a β(1-4)-Gal terminated (1-6) arm, as we have seen in the hybrid N-glycans constructs. Therefore, in β(1-2) xylosylated N-glycans terminating with β(1-3)-Gal, both arms should be more available for recognition, binding and further functionalization[30], unlike in mammalian N-glycans where the β(1-4)-Gal determines a prevalently closed and inaccessible (1-6) arm[24,27]. Also, the analysis of the structure and dynamics of the LeA terminating plant N-glycans showed that the specific branching and spatial orientation of the motif allowed for a stable interaction between the arms, which is not observed in complex N-glycans with a linear functionalization of the arms[24]. Notably, the same hydrogen bonding network between the arms cannot be established when the same N-glycan terminates with LeX, because of the non-complementary position of the α(1-3)-Fuc deoxy-C_6_.

The analysis of all these different complex N-glycoforms clearly shows that every modification, addition or removal of a specific motif, can greatly affect the 3D architecture of the N-glycan, thus its accessibility and complementarity to a receptor. However, these effects are rather complex to understand or to predict, if we think of the N-glycans 3D structure in terms of sequence of monosaccharides, a view that stems from the way we think about proteins. Our results show that the main effect of all functionalizations is actually local. For example, the core α(1-3)-Fuc forces a rotation of the chitobiose, a degree of freedom very lowly populated otherwise; meanwhile, β(1-2)-Xyl restricts the flexibility of the trimannose core and occupies its centre. Within this framework, the 3D structural and dynamics features of the N-glycoforms can be rationalized by discretizing their architecture in terms 3D units, or “glycoblocks”, that group monosaccharides and their linkages within their immediate spatial vicinity, e.g. the core α(1-3)-Fuc and the chitobiose which structure it has modified. A list of the glycoblocks that we have identified with the corresponding descriptors of their 3D features are listed in **Figure 7**. The whole N-glycan 3D architecture, in terms of the structures accessible and their conformational propensity, can be then described through the combination of these glycoblocks, together with the knowledge of their dynamic properties and flexibility. Also, consideration of these glycoblocks as spatial units can be useful to understand recognition by lectins and antibodies, which is often affected primarily by the targeted monosaccharide’s immediate vicinity and by its accessibility within a specific glycoform. For example, if we consider the 3D structure of the β(1-2)-Xyl Man3 glycoblock *vs*. the Man3 without Xyl, we can understand how the β(1-2)-Xyl position within that unit negates binding to DC-SIGN lectins[19], see **Figure S.3 panels a and b**. Additionally, we can see that the slight rotation on the chitobiose imposed by the core α(1-3)-Fuc does not prevent recognition and binding, see **Figure S.3 panel c**.

**Figure 7.**
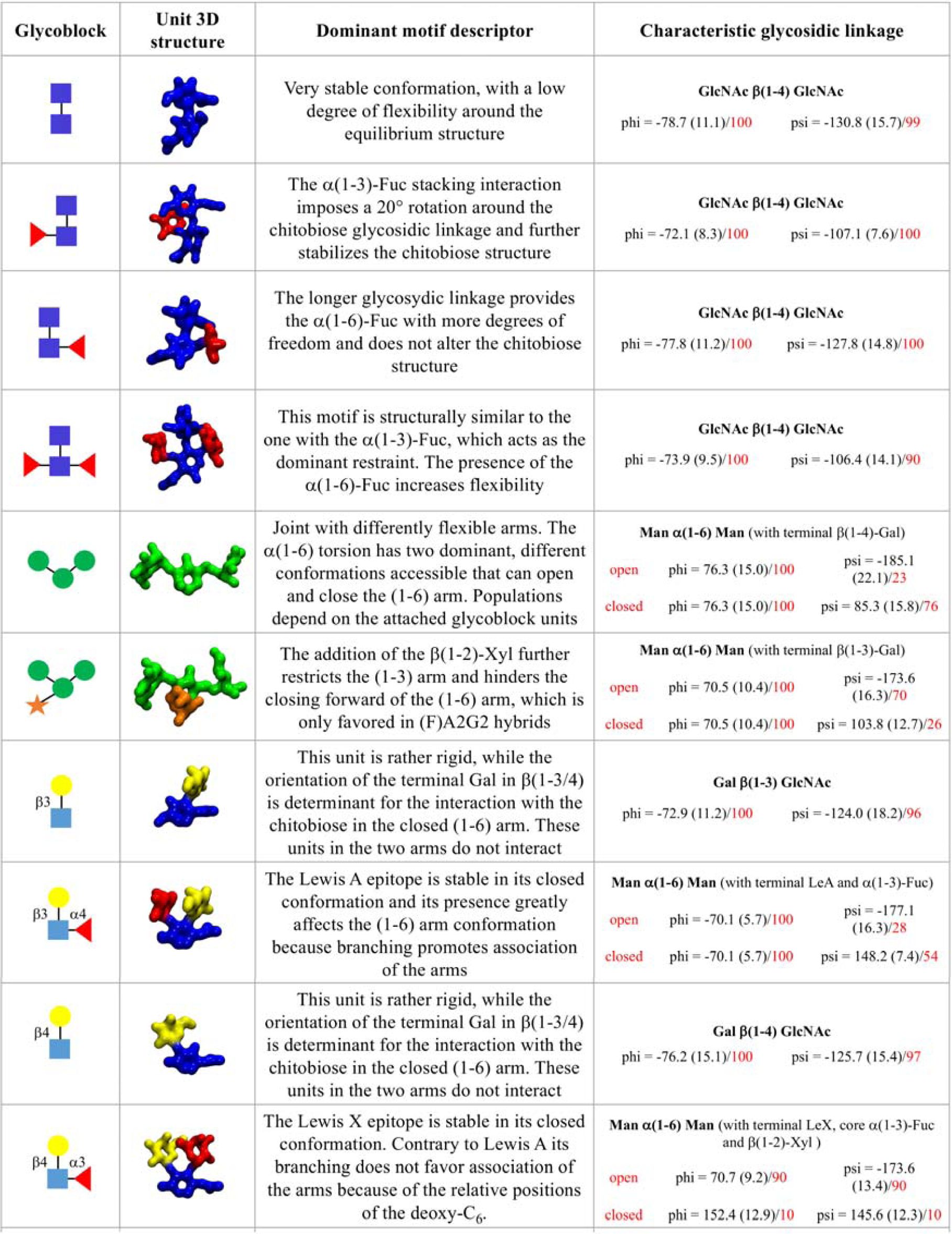
List of 3D structural units of monosaccharides (glycoblocks) that regulate the 3D architecture and dynamics of complex biantennary N-glycans from plants and invertebrate sources and hybrid mammalian constructs. The SFNG representation of each glycoblock is indicated in the first column from the left, 3D structures from the highest populated conformers are shown in the second column, rendered with VMD. A brief summary of the conformational features of each glycoblock and the characteristic linkage or its effect on the (1-6) arm conformation are indicates in the last two columns, respectively.

## Conclusions

In this work we used extensive sampling through MD simulations to study the effects on the N-glycan architecture of subtle, yet highly consequential modifications, namely core α(1-3)-Fuc and β(1-2)-Xyl[19]. These are part of standard N-glycoforms found in plants[23] and invertebrates[30], but immunogenic in humans[22,26,21]. Our results show that these modifications can greatly affect the 3D structure of the N-glycan and its structural dynamics, therefore its selective recognition by lectin receptors and antibodies. The atomistic-level of detail information that the MD simulations provide us with, highlights that the effects of different functionalizations, in terms of monosaccharide types and linkages, are primarily local, affecting the immediate spatial vicinity of the monosaccharide within the N-glycan structure. Within this framework, we propose an alternative approach that can help describe and predict the architecture of N-glycans based on the combination of structural 3D units, or glycoblocks. Unlike a description based on monosaccharide sequence and linkages as two separate features, the transition to well-defined and self-contained units, integrating information on both monosaccharides and linkages, can help us rationalize and deconvolute the glycans structural disorder and ultimately understand more clearly the relationships between sequence and structure in complex carbohydrates.

## Acknowledgements

The Irish Centre for High-End Computing (ICHEC) is gratefully acknowledged for generous allocation of computational resources. EF and CAF acknowledge the Irish Research Council (IRC) for funding through the Government of Ireland Postgraduate Scholarship Programme. EF and AMH acknowledge the John and Pat Hume Doctoral Scholarship Programme at Maynooth University for funding. EF would like to thank Prof. Iain B.H. Wilson for insightful feedback on an earlier version of the manuscript.

