## Supplementary Material for "How and why plants and human N-glycans are different: Insight from molecular dynamics into the “glycoblocks” architecture of complex carbohydrates"

**1. Computational Methods**

**System preparation.** All N-glycan starting structures for the MD simulations were generated with the GLYCAM Carbohydrate Builder (<http://www.glycam.org>). For each sequence we selected the complete set of rotamers obtained by variation of the 1-6 torsion angles, namely *gg*, *gt* and *tg* conformations for each 1-6 torsion. The topology file corresponding to each structure was obtained using *tleap^1^*, with parameters from the GLYCAM06-j1^2^ for the carbohydrate atoms and with TIP3P for water molecules^3^. Each N-glycan was placed in the centre of a cubic simulation box of 16 Å sides with no counterions to be consistent with the simulations run in earlier work^4^. Long range electrostatic were treated by Particle Mesh Ewald (PME) with cut-off set at 11 Å and a B-spline interpolation for mapping particles to and from the mesh of order of 4. Van der Waals (vdW) interactions were cut-off at 11 Å. The MD trajectories were generated by Langevin dynamics^5, 6^ with collision frequency of 1.0 ps^-1^. Pressure was kept constant by isotropic pressure scaling^7^ with a pressure relaxation time of 2.0 ps. Integration was done with a time step of 0.002 ps for all simulations, with bonds to hydrogen atoms restrained with the SHAKE algorithm^8^. All calculations were run with the AMBER18 software package^1^ on NVIDIA Tesla V100 16GB PCIe (Volta architecture) GPUs installed on the HPC infrastructure *kay* at the Irish Centre for High-End Computing (ICHEC).

**Simulation protocol.** The energy of the hydrated systems was initially minimized through 500,000 cycles of steepest descent, with all heavy atoms restrained with a harmonic potential with a force constant of 5 kcal mol^-1^Å^-2^. After minimization, the system was heated in two stages. During the first stage the temperature was raised from 0 to 100 K over 500 ps at constant volume and in the second stage from 100 K to 300 K over 500 ps at constant pressure. Through the heating process all heavy atoms were kept restrained. After heating phase all restraints were removed and the system was allowed to equilibrate for 5 ns at 300 K and at 1 atm of pressure. Separate production steps of 500 ns each were run for each rotamer (starting system) and convergence was assessed based on conformational and clustering analysis. Simulations were extended, if the sampling was not deemed as fully converged.

**Data analysis.** All trajectories were processed using *cpptraj^1^* and visually analysed with the Visual Molecular Dynamics (VMD) software package^9^. Backbone Root Mean Square Deviation (RMSD) and torsion angles values were measured using VMD. A density-based clustering method was used to calculate the populations of occupied conformations for each torsion angle in a trajectory and heat maps for each dihedral were generated with a kernel density estimate (KDE) function. Statistical and clustering analysis was done with the R package and data were plotted with RStudio ([www.rstudio.com](http://www.rstudio.com)).

**2. Supplementary Tables and Figures**

**Table S.1.** Results of the clustering analysis showing the median and standard deviation values (in parenthesis) for the torsion angles (°) measured through a cumulative 1.5 μs MD sampling of the α(1-3) core fucosylated ngf glycan.

| **Fuc α(1-3) GlcNAc** | **Phi** | **Psi** | **Pop(%)** |
| --- | --- | --- | --- |
| Cluster 1 | -71.1 (8.9) | 141.1 (6.3) | 100 |
| **GlcNAc β(1-4) GlcNAc** | **Phi** | **Psi** | **Pop(%)** |
| Cluster 1 | -72.1 (8.3) | -107.1 (7.6) | 100 |
| **Man β(1-4) GlcNAc** | **Phi** | **Psi** | **Pop(%)** |
| Cluster 1 | -73.3 (12.3) | -122.1 (14.9) | 90.0 |
| Cluster 2 | -166.4 (16.7) | -145.6 (9.8) | 8.3 |
| Cluster 3 | -64.0 (10.0) | 75.3 (9.3) | 1.7 |
| **Man α(1-6) Man** | **Phi** | **Psi** | **Pop(%)** |
| Cluster 1 | 69.2 (10.2) | -176.6 (17.5) | 65.7 |
| Cluster 2 | 74.2 (9.4) | 89.2 (10.8) | 25.2 |
| Cluster 3 | 60.4 (6.4) | 60.0 (5.8) | 9.1 |
| **GlcNAc β(1-2) Man (*1-6*)** | **Phi** | **Psi** | **Pop(%)** |
| Cluster 1 | -78.8 (14.3) | 160.7 (20.6) | 96.2 |
| Cluster 2 | 69.6 (9.3) | 154.8 (10.3) | 2.5 |
| **Man α(1-3) Man** | **Phi** | **Psi** | **Pop(%)** |
| Cluster 1 | 70.9 (8.5) | 140.3 (13.5) | 66.8 |
| Cluster 2 | 70.6 (8.7) | 99.2 (9.5) | 33.2 |
| **GlcNAc β(1-2) Man (*1-3*)** | **Phi** | **Psi** | **Pop(%)** |
| Cluster 1 | -77.8 (13.7) | 161.5 (13.5) | 88.8 |
| Cluster 2 | -78.5 (8.0) | 109.2 (7.2) | 9.7 |

**Table S.2.** Results of the clustering analysis showing the median and standard deviation values (in parenthesis) for the torsion angles (°) measured through a cumulative 1.5 μs MD sampling of the α(1-3) core fucosylated gf glycan.

| **Fuc α(1-3) GlcNAc** | **Phi** | **Psi** | **Pop(%)** |
| --- | --- | --- | --- |
| Cluster 1 | -72.5 (10.7) | 125.2 (17.4) | 100 |
| **GlcNAc β(1-4) GlcNAc** | **Phi** | **Psi** | **Pop(%)** |
| Cluster 1 | -72.9 (8.6) | -105.7 (11.7) | 100 |
| **Man β(1-4) GlcNAc** | **Phi** | **Psi** | **Pop(%)** |
| Cluster 1 | -74.0 (12.3) | -123.9 (15.7) | 88.5 |
| Cluster 2 | -165.7 (14.7) | -145.7 (8.8) | 11.5 |
| **Man α(1-6) Man** | **Phi** | **Psi** | **Pop(%)** |
| Cluster 1 | 69.3 (10.6) | -177.6 (19.4) | 72.5 |
| Cluster 2 | 71.5 (6.2) | 79.2 (7.1) | 16.2 |
| Cluster 3 | 60.7 (5.7) | 60.4 (5.1) | 1.1 |
| **GlcNAc β(1-2) Man (*1-6*)** | **Phi** | **Psi** | **Pop(%)** |
| Cluster 1 | -76.7 (13.8) | 162.5 (12.4) | 88.4 |
| Cluster 2 | -76.0 (6.2) | 113.5 (6.6) | 5.9 |
| Cluster 3 | -147.6 (9.5) | 97.9 (9.4) | 1.9 |
| **Gal β(1-3) GlcNAc (*1-6*)** | **Phi** | **Psi** | **Pop(%)** |
| Cluster 1 | -72.9 (11.2) | 124.0 (18.2) | 96.6 |
| Cluster 2 | -82.1 (12.2) | -64.0 (8.4) | 3.0 |
| Cluster 3 | -150.0 (11.3) | -101.0 (7.3) | 1.4 |
| **Man α(1-3) Man** | **Phi** | **Psi** | **Pop(%)** |
| Cluster 1 | 71.7 (8.9) | 141.5 (14.5) | 66.9 |
| Cluster 2 | 70.4 (8.6) | 99.12 (9.5) | 33.1 |
| **GlcNAc β(1-2) Man (*1-3*)** | **Phi** | **Psi** | **Pop(%)** |
| Cluster 1 | -77.9 (14.2) | 160.9 (14.5) | 83.3 |
| Cluster 2 | 67.2 (9.2) | 152.4 (10.4) | 8.6 |
| Cluster 3 | -77.7 (7.7) | 152.4 (6.2) | 8.1 |
| **Gal β(1-3) GlcNAc (*1-3*)** | **Phi** | **Psi** | **Pop(%)** |
| Cluster 1 | -72.4 (9.8) | 125.0 (15.8) | 100 |

**Table S.3.** Results of the clustering analysis showing the median and standard deviation values (in parenthesis) for the torsion angles (°) measured through a cumulative 3 μs MD sampling of the α(1-2) core xylosylated ngx glycan.

| **GlcNAc β(1-4) GlcNAc** | **Phi** | **Psi** | **Pop(%)** |
| --- | --- | --- | --- |
| Cluster 1 | -78.1 (10.2) | -129.6 (15.67) | 96.3 |
| Cluster 2 | -81.8 (9.8) | 63.8 (8.11) | 3.7 |
| **Man β(1-4) GlcNAc** | **Phi** | **Psi** | **Pop(%)** |
| Cluster 1 | -75.5 (12.6) | -123.5 (14.6) | 94.4 |
| Cluster 2 | -64.2 (6.8) | 74.2 (9.7) | 5.6 |
| **Xyl β(1-2) Man** | **Phi** | **Psi** | **Pop(%)** |
| Cluster 1 | -80.2 (12.0) | 133.2 (17.0) | 100 |
| **Man α(1-6) Man** | **Phi** | **Psi** | **Pop(%)** |
| Cluster 1 | 69.7 (10.3) | -174.4 (16.8) | 77.2 |
| Cluster 2 | 73.5 (10.1) | 106.1 (10.6) | 22.8 |
| **GlcNAc β(1-2) Man (*1-6*)** | **Phi** | **Psi** | **Pop(%)** |
| Cluster 1 | -80.9 (15.5) | 161.7 (12.0) | 90.1 |
| Cluster 2 | -73.1 (7.9) | 114.2 (7.8) | 9.3 |
| **Man α(1-3) Man** | **Phi** | **Psi** | **Pop(%)** |
| Cluster 1 | 78.4 (7.3) | 114.8 (14.8) | 100 |
| **GlcNAc β(1-2) Man (*1-3*)** | **Phi** | **Psi** | **Pop(%)** |
| Cluster 1 | -77.4 (13.3) | 161.5 (12.2) | 88.2 |
| Cluster 2 | -78.9 (6.4) | 109.1 (7.18) | 8.2 |
| Cluster 3 | -66.9 (9.0) | 149.1 (12.3) | 3.6 |

**Table S.4.** Results of the clustering analysis showing the median and standard deviation values (in parenthesis) for the torsion angles (°) measured through a cumulative 1.5 μs MD sampling of the α(1-3) core xylosylated gx glycan.

| **GlcNAc β(1-4) GlcNAc** | **Phi** | **Psi** | **Pop(%)** |
| --- | --- | --- | --- |
| Cluster 1 | -78.0 (10.1) | -130.2(15.7) | 97.5 |
| Cluster 2 | -84.0 (5.4) | -64.8 (5.6) | 2.5 |
| **Man β(1-4) GlcNAc** | **Phi** | **Psi** | **Pop(%)** |
| Cluster 1 | -75.2 (13.5) | -124.7 (15.0) | 87.6 |
| Cluster 2 | -67.9 (14.7) | 72.9 (11.7) | 11.3 |
| Cluster 3 | -178.3 (6.9) | -175.5 (7.3) | 1.0 |
| **Xyl β(1-2) Man** | **Phi** | **Psi** | **Pop(%)** |
| Cluster 1 | -78.2 (9.3) | 138.4 (15.4) | 100 |
| **Man α(1-6) Man** | **Phi** | **Psi** | **Pop(%)** |
| Cluster 1 | 70.5 (10.4) | -173.6 (19.4) | 70.0 |
| Cluster 2 | 71.8 (9.6) | 103.8 (12.9) | 25.7 |
| Cluster 3 | 161.6 (8.0) | 132.5 (9.2) | 2.3 |
| Cluster 4 | 79.0 (6.7) | -80.2 (8.05) | 1.59 |
| **GlcNAc β(1-2) Man (*1-6*)** | **Phi** | **Psi** | **Pop(%)** |
| Cluster 1 | -82.1 (15.0) | 161.3 (12.2) | 88.2 |
| Cluster 2 | -77.7 (7.47) | 112.5 (7.0) | 10.0 |
| Cluster 3 | -147.3 (9.0) | 98.9 (9.4) | 1.9 |
| **Gal β(1-3) GlcNAc (*1-6*)** | **Phi** | **Psi** | **Pop(%)** |
| Cluster 1 | -72.1 (10.0) | 126.1 (15.5) | 80.8 |
| Cluster 2 | -83.6 (11.4) | -63.0 (7.57) | 19.2 |
| **Man α(1-3) Man** | **Phi** | **Psi** | **Pop(%)** |
| Cluster 1 | 72.5 (10.7) | 125.2 (17.4) | 100 |
| **GlcNAc β(1-2) Man (*1-3*)** | **Phi** | **Psi** | **Pop(%)** |
| Cluster 1 | -78.5 (14.2) | 162.4 (13.0) | 91.2 |
| Cluster 2 | -77.8 (7.4) | 111.1 (7.0) | 8.8 |
| **Gal β(1-3) GlcNAc (*1-3*)** | **Phi** | **Psi** | **Pop(%)** |
| Cluster 1 | -71.1 (7.2) | 125.5 (11.8) | 100 |

**Table S.5.** Results of the clustering analysis showing the median and standard deviation values (in parenthesis) for the torsion angles (°) measured through a cumulative 1.5 μs MD sampling of the β(1-2) core xylosylated and α(1-3) core fucosylated ngxf glycan.

| **Fuc α(1-3) GlcNAc** | **Phi** | **Psi** | **Pop(%)** |
| --- | --- | --- | --- |
| Cluster 1 | -71.3 (9.0) | 140.5 (6.0) | 100 |
| **GlcNAc β(1-4) GlcNAc** | **Phi** | **Psi** | **Pop(%)** |
| Cluster 1 | -73.2 (11.0) | -106.4 (21.0) | 100 |
| **Man β(1-4) GlcNAc** | **Phi** | **Psi** | **Pop(%)** |
| Cluster 1 | -75.9 (14.4) | -124.2 (15.3) | 75.4 |
| Cluster 2 | -66.5 (11.3) | 73.3 (10.8) | 21.1 |
| Cluster 3 | 179.0 (7.6) | -174.7 (8.2) | 2.3 |
| Cluster 4 | -151.9 (8.2) | -146.5 (6.7) | 1.2 |
| **Xyl β(1-2) Man** | **Phi** | **Psi** | **Pop(%)** |
| Cluster 1 | -78.8 (10.8) | 135.9 (17.4) | 100 |
| **Man α(1-6) Man** | **Phi** | **Psi** | **Pop(%)** |
| Cluster 1 | 70.6 (9.0) | 175.2 (17.6) | 68.5 |
| Cluster 2 | 71.7 (10.0) | 106.5 (12.3) | 24.5 |
| Cluster 3 | 81.6 (8.0) | -75.0 (10.0) | 4.4 |
| Cluster 4 | 158.0 (8.9) | 135.9 (10.4) | 2.4 |
| **GlcNAc β(1-2) Man (*1-6*)** | **Phi** | **Psi** | **Pop(%)** |
| Cluster 1 | -79.2 (14.4) | 158.4 (24.9) | 90.1 |
| Cluster 2 | -80.0 (6.33) | 113.0 (7.3) | 9.3 |
| **Man α(1-3) Man** | **Phi** | **Psi** | **Pop(%)** |
| Cluster 1 | 69.1 (9.4) | 113.8 (16.8) | 1 |
| **GlcNAc β(1-2) Man (*1-3*)** | **Phi** | **Psi** | **Pop(%)** |
| Cluster 1 | -77.4 (13.3) | 161.5 (12.2) | 88.2 |
| Cluster 2 | -78.9 (6.4) | 109.1 (7.18) | 8.21 |
| Cluster 3 | -66.9 (9.0) | 149.1 (12.3) | 3.58 |

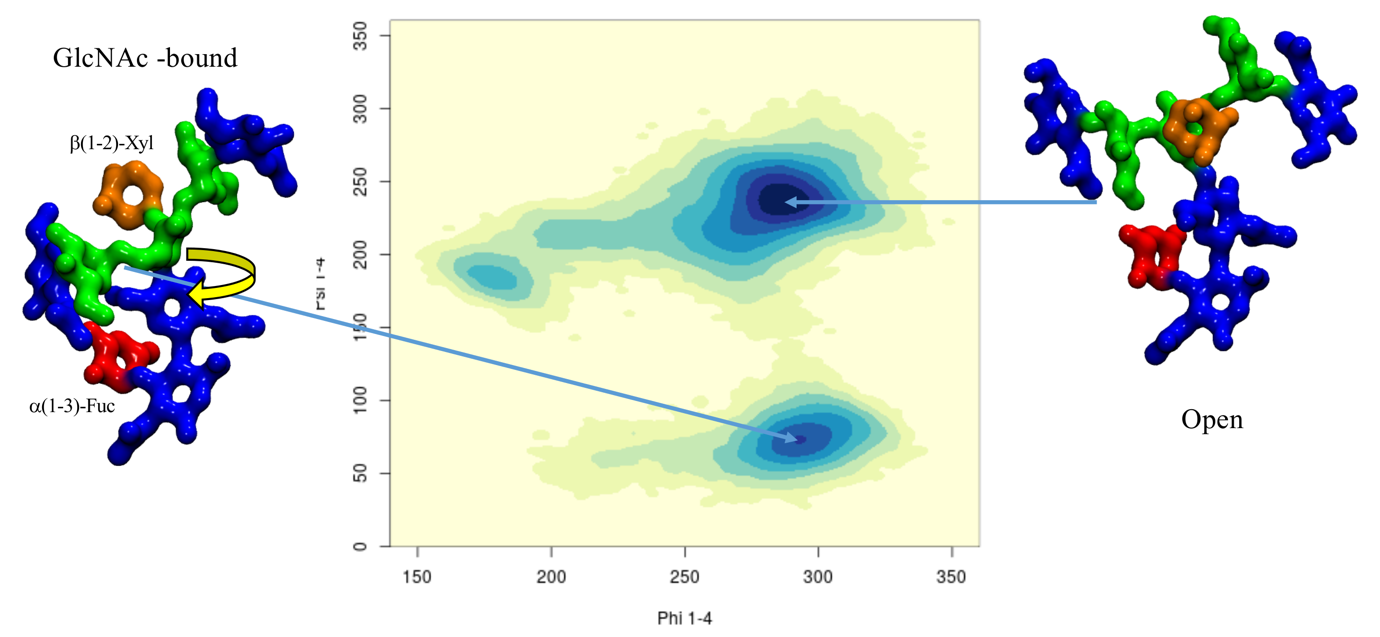

**Figure S.1.** Conformational analysis of the ngxf glycoform in terms of phi/psi torsion values, shown on the x and y axis, respectively, of the core Man-β(1-4)-GlcNAc linkage. Representative structures selected from MD sampling are shown on the left- and right-hand side of the heat map. The GlcNAc-bound conformation is obtained through a torsion of the Man3 “glycoblock” relative to the chitobiose and it is stabilized by hydrogen bonding interactions between the β(1-2)-Xyl and α(1-3)-Fuc. The monosaccharides colouring follows the SFNG nomenclature. The structure rendering was done with VMD and the graphical statistical analysis with RStudio ([www.rstudio.com](http://www.rstudio.com)).

**Table S.6** Results of the clustering analysis showing the median and standard deviation values (in parenthesis) for the torsion angles (°) measured through a cumulative 1.5 μs MD sampling of the β(1-2) xylosylated and α(1-3) core fucosylated gxf glycan

| **Fuc α(1-3) GlcNAc** | **Phi** | **Psi** | **Pop(%)** |
| --- | --- | --- | --- |
| Cluster 1 | -71.1 (9.8) | 140.5 (7.1) | 88.9 |
| Cluster 2 | -156.8 (5.6) | 90.7 (6.7) | 11.1 |
| **GlcNAc β(1-4) GlcNAc** | **Phi** | **Psi** | **Pop(%)** |
| Cluster 1 | -72.2 (8.5) | -107.0 (8.0) | 100 |
| **Man β(1-4) GlcNAc** | **Phi** | **Psi** | **Pop(%)** |
| Cluster 1 | -76.7 (15.1) | -124.5 (17.0) | 85.2 |
| Cluster 2 | -179.5 (8.5) | -174.7 (9.5) | 8.3 |
| Cluster 3 | -63.0 (9.9) | -75.6 (10.3) | 5.5 |
| Cluster 4 | -153.1 (6.2) | -147.4 (5.31) | 1 |
| **Xyl β(1-2) Man** | **Phi** | **Psi** | **Pop(%)** |
| Cluster 1 | -78.2 (9.9) | 135.9 (17.4) | 100 |
| **Man α(1-6) Man** | **Phi** | **Psi** | **Pop(%)** |
| Cluster 1 | 70.73 (10.3) | -174.2 (19.0) | 87.2 |
| Cluster 2 | 70.8 (8.1) | 101.5 (10.7) | 9.3 |
| Cluster 3 | 161.6 (8.5) | 131.9 (10.6) | 3.5 |
| **GlcNAc β(1-2) Man (*1-6*)** | **Phi** | **Psi** | **Pop(%)** |
| Cluster 1 | -78.1 (14.3) | 161.9 (12.9) | 84.7 |
| Cluster 2 | -76.1 (7.6) | 111.6 (7.2) | 10.5 |
| Cluster 3 | -147.1 (10.9) | 99.5 (10.1) | 3.2 |
| Cluster 4 | 67.4 (7.4) | 152.4 (8.4) | 1.6 |
| **Gal β(1-3) GlcNAc (*1-6*)** | **Phi** | **Psi** | **Pop(%)** |
| Cluster 1 | -70.7 (10.4) | 125.0 (16.8) | 96.7 |
| Cluster 2 | -81.6 (7.07) | -64.6 (6.4) | 3.3 |
| Cluster 3 | -81.6 (7.07) | -64.6 (6.4) | 1.3 |
| **Man α(1-3) Man** | **Phi** | **Psi** | **Pop(%)** |
| Cluster 1 | 68.7 (9.1) | 114.1 (16.6) | 1 |
| **GlcNAc β(1-2) Man (*1-3*)** | **Phi** | **Psi** | **Pop(%)** |
| Cluster 1 | -78.2 (14.7) | 162.0 (13.0) | 90.1 |
| Cluster 2 | -78.1 (7.2) | 112.1 (6.8) | 9.9 |
| **Gal β(1-3) GlcNAc (*1-3*)** | **Phi** | **Psi** | **Pop(%)** |
| Cluster 1 | -71.9 (6.9) | 126.0 (11.9) | 100 |

**Table S.7.** Results of the clustering analysis showing the median and standard deviation values (in parenthesis) for the torsion angles (°) measured through a cumulative 1.5 μs MD sampling of the α(1-3) core fucosylated LeA glycan.

| **Fuc α(1-3) GlcNAc** | **sPhi** | **Psi** | **Pop(%)** |
| --- | --- | --- | --- |
| Cluster 1 | -71.7 (9.8) | 141.8 (6.6) | 82.8 |
| Cluster 2 | -156.3 (7.02) | 88.8 (17.6) | 17.2 |
| **GlcNAc β(1-4) GlcNAc** | **Phi** | **Psi** | **Pop(%)** |
| Cluster 1 | -72.6 (8.7) | -107.4 (7.4) | 100 |
| **Man β(1-4) GlcNAc** | **Phi** | **Psi** | **Pop(%)** |
| Cluster 1 | -72.2 (12.0) | -124.5 (14.0) | 97.1 |
| Cluster 2 | 179.7 (6.1) | -178.2 (6.6) | 2.9 |
| **Man α(1-6) Man** | **Phi** | **Psi** | **Pop(%)** |
| Cluster 1 | -75.5 (7.4) | 148.1 (7.4) | 53.6 |
| Cluster 2 | -70.1 (5.8) | -177.0 (11.2) | 28.2 |
| Cluster 3 | -73.2 (6.7) | -101.9 (6.0) | 9.4 |
| Cluster 4 | -148.0 (6.3) | -165.3 (5.8) | 8.8 |
| **Xyl β(1-2) Man** | **Phi** | **Psi** | **Pop(%)** |
| Cluster 1 | -83.9 (10.5) | 132.8 (14.0) | 100 |
| **GlcNAc β(1-2) Man (*1-6*)** | **Phi** | **Psi** | **Pop(%)** |
| Cluster 1 | -150.0 (9.3) | 98.5 (7.9) | 40.9 |
| Cluster 2 | -91.5 (8.5) | 151.6 (8.6) | 29.9 |
| Cluster 3 | -62.7 (8.5) | 161.4 (9.9) | 29.2 |
| **Gal β(1-3) GlcNAc (*1-6*)** | **Phi** | **Psi** | **Pop(%)** |
| Cluster 1 | -70.1 (7.6) | 131.5 (7.6) | 100 |
| **Fuc α(1-4) GlcNAc (*1-6*)** | **Phi** | **Psi** | **Pop(%)** |
| Cluster 1 | -67.8 (7.5) | -101.26 (7.3) | 100 |
| **Man α(1-3) Man** | **Phi** | **Psi** | **Pop(%)** |
| Cluster 1 | 61.5 (8.5) | 111.0 (15.1) | 100 |
| **GlcNAc β(1-2) Man (*1-3*)** | **Phi** | **Psi** | **Pop(%)** |
| Cluster 1 | -77.9 (16.9) | 162.3 (12.3) | 57.8 |
| Cluster 2 | -79.9 (11.4) | 105.1 (12.4) | 27.9 |
| Cluster 3 | 66.1 (10.1) | 153.8 (10.8) | 14.26 |
| **Gal β(1-3) GlcNAc (*1-3*)** | **Phi** | **Psi** | **Pop(%)** |
| Cluster 1 | -70.6 (7.5) | 134.5 (6.8) | 100 |
| **Fuc α(1-4) GlcNAc (*1-3*)** | **Phi** | **Psi** | **Pop(%)** |
| Cluster 1 | -68.6 (8.9) | -100.9 (6.7) | 98.4 |
| Cluster 2 | -148.9 (8.0) | -150.7 (3.8) | 1.6 |

**Table S.8** Results of the clustering analysis showing the median and standard deviation values (in parenthesis) for the torsion angles (°) measured through a cumulative 1.5 μs MD sampling of the β(1-2) xylosylated mgx glycan. Note: mg refers to the mammalian terminal β(1-4)-Gal.

| **GlcNAc β(1-4) GlcNAc** | **Phi** | **Psi** | **Pop(%)** |
| --- | --- | --- | --- |
| Cluster 1 | -78.2 (10.9) | -131.1 (15.8) | 97.5 |
| Cluster 2 | -79.6 (11.3) | 66.6 (11.5) | 2.5 |
| **Man β(1-4) GlcNAc** | **Phi** | **Psi** | **Pop(%)** |
| Cluster 1 | -75.7 (17.1) | -123.7 (14.7) | 91.3 |
| Cluster 2 | -68.1 (12.6) | 72.1 (11.9) | 8.7 |
| **Xyl β(1-2) Man** | **Phi** | **Psi** | **Pop(%)** |
| Cluster 1 | -81.7 (19.1) | 133.6 (20.1) | 100 |
| **Man α(1-6) Man** | **Phi** | **Psi** | **Pop(%)** |
| Cluster 1 | 72.2 (9.4) | 103.5 (11.3) | 56.9 |
| Cluster 2 | 69.9 (8.3) | -173.8 (15.7) | 39.8 |
| Cluster 3 | 162.2 (9.1) | 131.2 (9.3) | 1.7 |
| Cluster 4 | 58.5 (3.2) | 59.98 (4.3) | 1.6 |
| **GlcNAc β(1-2) Man (*1-6*)** | **Phi** | **Psi** | **Pop(%)** |
| Cluster 1 | -78.1 (14.3) | 161.9 (12.9) | 84.7 |
| Cluster 2 | -76.1 (7.6) | 111.6 (7.2) | 10.5 |
| Cluster 3 | -147.1 (10.9) | 99.5 (10.1) | 3.2 |
| Cluster 4 | 67.4 (7.4) | 152.4 (8.4) | 1.6 |
| **Gal β(1-4) GlcNAc (*1-6*)** | **Phi** | **Psi** | **Pop(%)** |
| Cluster 1 | -72.9 (15.8) | -119.9 (16.0) | 98.9 |
| Cluster 2 | -73.5 (12.6) | -73.24 (12.5) | 0.6 |
| Cluster 3 | 63.6 (10.7) | -117.8 (6.8) | 0.8 |
| **Man α(1-3) Man** | **Phi** | **Psi** | **Pop(%)** |
| Cluster 1 | 68.9 (9.7) | 114.5 (16.7) | 1 |
| **GlcNAc β(1-2) Man (*1-3*)** | **Phi** | **Psi** | **Pop(%)** |
| Cluster 1 | -79.0 (13.8) | 162.5 (12.2) | 87.5 |
| Cluster 2 | -80.9 (9.12) | 109.4 (9.0) | 12.5 |
| **Gal β(1-4) GlcNAc (*1-3*)** | **Phi** | **Psi** | **Pop(%)** |
| Cluster 1 | -72.41 (10.9) | -118.4 (15.1) | 100 |

**Table S.9** Results of the clustering analysis showing the median and standard deviation values (in parenthesis) for the torsion angles (°) measured through a cumulative 4.5 μs MD sampling of the β(1-2) xylosylated and α(1-6) core fucosylated mgmfx glycan. Note: mg refers to the mammalian terminal β(1-4)-Gal and mf to the mammalian core α(1-6)-Fuc.

| **Fuc α(1-6) GlcNAc** | **Phi** | **Psi** | **Pop(%)** |
| --- | --- | --- | --- |
| Cluster 1 | -74.5 (9.6) | 172.4 (14.5) | 92.1 |
| Cluster 2 | -95.9 (4.5) | 71.78 (6.1) | 6.9 |
| Cluster 3 | -75.6 (2.5) | 1113.8 (2.17) | 1.0 |
| **GlcNAc β(1-4) GlcNAc** | **Phi** | **Psi** | **Pop(%)** |
| Cluster 1 | -77.2 (9.5) | -126.0 (14.3) | 100 |
| **Man β(1-4) GlcNAc** | **Phi** | **Psi** | **Pop(%)** |
| Cluster 1 | -74.3 (15.0) | -122.7 (14.2) | 97.1 |
| Cluster 2 | -67.4 (10.1) | 73.6 (10.7) | 2.9 |
| **Man α(1-6) Man** | **Phi** | **Psi** | **Pop(%)** |
| Cluster 1 | 71.8 (9.4) | 103.3 (10.98) | 70.0 |
| Cluster 2 | 70.4 (9.0) | -177.0 (11.2) | 26.9 |
| Cluster 3 | 67.5 (5.24) | -62.5 (5.7) | 3.1 |
| **Xyl β(1-2) Man** | **Phi** | **Psi** | **Pop(%)** |
| Cluster 1 | -83.9 (10.5) | 132.8 (14.0) | 100 |
| **GlcNAc β(1-2) Man (*1-6*)** | **Phi** | **Psi** | **Pop(%)** |
| Cluster 1 | -91.5 (12.5) | 159.9 (10.3) | 96.3 |
| Cluster 2 | -75.5 (3.9) | 113.6 (3.8) | 2.9 |
| Cluster 3 | 66.0 (6.8) | 154.3 (5.8) | 0.8 |
| **Gal β(1-4) GlcNAc (*1-6*)** | **Phi** | **Psi** | **Pop(%)** |
| Cluster 1 | -74.9 (13.56) | -122.1 (15.5) | 97.5 |
| Cluster 2 | -83.6 (18.6) | 65.3 (13.5) | 2.5 |
| **Man α(1-3) Man** | **Phi** | **Psi** | **Pop(%)** |
| Cluster 1 | 68.7 (9.9) | 115.6 (16.9) | 100 |
| **GlcNAc β(1-2) Man (*1-3*)** | **Phi** | **Psi** | **Pop(%)** |
| Cluster 1 | -79.1 (13.2) | 162.3 (11.4) | 90.9 |
| Cluster 2 | -77.9 (6.8) | 110.4 (7.6) | 9.1 |
| **Gal β(1-4) GlcNAc (*1-3*)** | **Phi** | **Psi** | **Pop(%)** |
| Cluster 1 | -72.7 (15.4) | -118.6 (15.7) | 97.3 |
| Cluster 2 | -74.0 (12.4) | 70.12 (13.1) | 2.7 |

**Table S.10** Results of the clustering analysis showing the median and standard deviation values (in parenthesis) for the torsion angles (°) measured through a cumulative 4.5 μs MD sampling of the β(1-2) xylosylated and α(1-3) core fucosylated nmgmfx glycan. Note: nmg refers to the absence of mammalian terminal β(1-4)-Gal and mf to the mammalian core α(1-6)-Fuc.

| **Fuc α(1-6) GlcNAc** | **Phi** | **Psi** | **Pop(%)** |
| --- | --- | --- | --- |
| Cluster 1 | -73.0 (9.6) | 177.0 (14.8) | 93.1 |
| Cluster 2 | -95.6 (4.0) | 75.0 (5.4) | 6.9 |
| **GlcNAc β(1-4) GlcNAc** | **Phi** | **Psi** | **Pop(%)** |
| Cluster 1 | -77.2 3(9.7) | -126.0 (14.4) | 100 |
| **Man β(1-4) GlcNAc** | **Phi** | **Psi** | **Pop(%)** |
| Cluster 1 | -76.6 (18.5) | -124.8 (16.9) | 91.1 |
| Cluster 2 | -66.2 (12.5) | 73.0 (11.9) | 8.2 |
| **Man α(1-6) Man** | **Phi** | **Psi** | **Pop(%)** |
| Cluster 1 | 70.1 (9.2) | -173.6 (14.6) | 71.5 |
| Cluster 2 | 72.1 (8.3) | 104.25 (9.7) | 28.5 |
| **Xyl β(1-2) Man** | **Phi** | **Psi** | **Pop(%)** |
| Cluster 1 | -80.6 (16.5) | 135.8 (14.0) | 100 |
| **GlcNAc β(1-2) Man (*1-6*)** | **Phi** | **Psi** | **Pop(%)** |
| Cluster 1 | -82.8 (14.94) | 161.1 (11.9) | 90.6 |
| Cluster 2 | -77.8 (7.3) | 111.3 (6.8) | 7.4 |
| Cluster 3 | 66.0 (7.3) | 152.9 (8.7) | 2.0 |
| **Man α(1-3) Man** | **Phi** | **Psi** | **Pop(%)** |
| Cluster 1 | 68.9 (9.4) | 114.4 (16.9) | 100 |
| **GlcNAc β(1-2) Man (*1-3*)** | **Phi** | **Psi** | **Pop(%)** |
| Cluster 1 | -78.4 (13.4) | 162.6 (11.4) | 88.8 |
| Cluster 2 | -78.9 (7.2) | 110.4 (7.0) | 9.0 |
| Cluster 2 | 66.45 (10.9) | 152.6 (10.7) | 2.2 |

**Table S.11** Results of the clustering analysis showing the median and standard deviation values (in parenthesis) for the torsion angles (°) measured through a cumulative 1.5 μs MD sampling of the β(1-2) xylosylated and α(1-3) core fucosylated mgpfx glycan. Note: mg refers to the mammalian terminal β(1-4)-Gal and pf to the plant core α(1-3)-Fuc.

| **Fuc α(1-3) GlcNAc** | **Phi** | **Psi** | **Pop(%)** |
| --- | --- | --- | --- |
| Cluster 1 | -70.8 (10.7) | 141.3 (8.9) | 93.8 |
| Cluster 2 | -156.8 (7.7) | 91.3 (9.3) | 6.2 |
| **GlcNAc β(1-4) GlcNAc** | **Phi** | **Psi** | **Pop(%)** |
| Cluster 1 | -72.5 (8.8) | -107.3 (9.1) | 100 |
| **Man β(1-4) GlcNAc** | **Phi** | **Psi** | **Pop(%)** |
| Cluster 1 | -76.4 (15.1) | -123.6 (17.3) | 87.0 |
| Cluster 2 | -68.2 (12.8) | 70.9 (11.2) | 13.0 |
| **Xyl β(1-2) Man** | **Phi** | **Psi** | **Pop(%)** |
| Cluster 1 | -80.7 (15.3) | 134.5 (16.5) | 100 |
| **Man α(1-6) Man** | **Phi** | **Psi** | **Pop(%)** |
| Cluster 1 | 69.9 (9.0) | -175.5 (15.0) | 49.1 |
| Cluster 2 | 72.4 (9.7) | 105.1 (12.5) | 46.7 |
| Cluster 3 | 158.6 (11.8) | 135.8 (13.6) | 4.2 |
| **GlcNAc β(1-2) Man (*1-6*)** | **Phi** | **Psi** | **Pop(%)** |
| Cluster 1 | -86.2 (15.0) | 160.9 (11.4) | 96.4 |
| Cluster 2 | -78.5 (6.1) | 113.5 (4.9) | 3.6 |
| **Gal β(1-4) GlcNAc (*1-6*)** | **Phi** | **Psi** | **Pop(%)** |
| Cluster 1 | -72.9 (15.0) | -119.9 (15.6) | 97.2 |
| Cluster 2 | -74.1 (13.1) | 70.1 (13.8) | 2.8 |
| **Man α(1-3) Man** | **Phi** | **Psi** | **Pop(%)** |
| Cluster 1 | 68.8 (9.6) | 114.5 (16.7) | 1 |
| **GlcNAc β(1-2) Man (*1-3*)** | **Phi** | **Psi** | **Pop(%)** |
| Cluster 1 | -78.8 (13.8) | 162.6 (12.6) | 88.3 |
| Cluster 2 | -80.7 (8.0) | 109.5 (7.6) | 9.6 |
| Cluster 3 | 65.2 (9.9) | 150.8 (10.8) | 2.1 |
| **Gal β(1-4) GlcNAc (*1-3*)** | **Phi** | **Psi** | **Pop(%)** |
| Cluster 1 | -72.5 (11.5) | -118.6 (15.5) | 100 |

**Table S.12** Results of the clustering analysis showing the median and standard deviation values (in parenthesis) for the torsion angles (°) measured through a cumulative 1.5 μs MD sampling of the α(1-3) core fucosylated mgpf glycan. Note: mg refers to the mammalian terminal β(1-4)-Gal and pf to the plant core α(1-3)-Fuc.

| **Fuc α(1-3) GlcNAc** | **Phi** | **Psi** | **Pop(%)** |
| --- | --- | --- | --- |
| Cluster 1 | -71.1 (10.6) | 140.3 (14.8) | 98.1 |
| Cluster 2 | -156.5 (6.2) | 91.4 (9.3) | 1.9 |
| **GlcNAc β(1-4) GlcNAc** | **Phi** | **Psi** | **Pop(%)** |
| Cluster 1 | -73.0 (14.8) | -121.5 (15.7) | 95.9 |
| Cluster 2 | -80.9 (15.3) | 62.5 (13.4) | 4.1 |
| **Man β(1-4) GlcNAc** | **Phi** | **Psi** | **Pop(%)** |
| Cluster 1 | -76.6 (14.1) | -124.9 (16.1) | 77.9 |
| Cluster 2 | -153.7 (13.3) | -139.7 (8.5) | 12.8 |
| Cluster 3 | -71.1 (12.5) | 69.6 (11.4) | 9.3 |
| **Man α(1-6) Man** | **Phi** | **Psi** | **Pop(%)** |
| Cluster 1 | 74.2 (13.0) | 86.5 (14.7) | 74.8 |
| Cluster 2 | 70.3 (9.1) | -176.5 (14.7) | 25.2 |
| **GlcNAc β(1-2) Man (*1-6*)** | **Phi** | **Psi** | **Pop(%)** |
| Cluster 1 | -80.2 (14.6) | 163.2 (12.4) | 100 |
| **Gal β(1-4) GlcNAc (*1-6*)** | **Phi** | **Psi** | **Pop(%)** |
| Cluster 1 | -73.0 (14.88) | -121.58 (15.8) | 95.9 |
| Cluster 2 | -80.9 (15.3) | 62.5 (13.4) | 4.1 |
| **Man α(1-3) Man** | **Phi** | **Psi** | **Pop(%)** |
| Cluster 1 | 70.95 (9.6) | 140.95 (15.2) | 73.0 |
| Cluster 1 | 70.1(8.21) | 101.2 (8.8) | 27.0 |
| **GlcNAc β(1-2) Man (*1-3*)** | **Phi** | **Psi** | **Pop(%)** |
| Cluster 1 | -78.5 (13.3) | 162.1 (11.7) | 91.2 |
| Cluster 2 | -77.9 (6.4) | 110.7 (6.62) | 8.8 |
| **Gal β(1-4) GlcNAc (*1-3*)** | **Phi** | **Psi** | **Pop(%)** |
| Cluster 1 | -72.2 (10.9) | -118.4 (15.0) | 100 |

**Table S.13** Results of the clustering analysis showing the median and standard deviation values (in parenthesis) for the torsion angles (°) measured through a cumulative 2 μs MD sampling of the α(1-3) and α(1-6) core fucosylated mgmfpf glycan. Note: mg refers to the mammalian terminal β(1-4)-Gal, pf to the plant core α(1-3)-Fuc and mf to the mammalian core α(1-6)-Fuc.

| **Fuc α(1-6) GlcNAc** | **Phi** | **Psi** | **Pop(%)** |
| --- | --- | --- | --- |
| Cluster 1 | -72.0 (9.6) | -179.5 (14.8) | 74.3 |
| Cluster 2 | -76.2 (4.0) | 117.36 (12.1) | 12.9 |
| Cluster 3 | -144.4 (7.7) | 171.0 (5.5) | 12.8 |
| **Fuc α(1-3) GlcNAc** | **Phi** | **Psi** | **Pop(%)** |
| Cluster 1 | -70.2 (10.3) | 140.1 (9.7) | 88.6 |
| Cluster 2 | -157.3 (7.7) | 91.4 (8.9) | 11.4 |
| **GlcNAc β(1-4) GlcNAc** | **Phi** | **Psi** | **Pop(%)** |
| Cluster 1 | -73.8 (9.5) | -106.4 (14.1) | 91.7 |
| Cluster 2 | -154.9 (10.6) | -147.8 (7.4) | 8.3 |
| **Man β(1-4) GlcNAc** | **Phi** | **Psi** | **Pop(%)** |
| Cluster 1 | -72.1 (10.9) | -120.7 (12.6) | 74.8 |
| Cluster 2 | -153.0 (13.0) | -139.9 (8.3) | 25.2 |
| **Man α(1-6) Man** | **Phi** | **Psi** | **Pop(%)** |
| Cluster 1 | 73.5 (11.7) | 86.6 (17.32) | 85.1 |
| Cluster 2 | 69.5 (9.9) | -176.3 (15.8) | 14.9 |
| **GlcNAc β(1-2) Man (*1-6*)** | **Phi** | **Psi** | **Pop(%)** |
| Cluster 1 | -85.1 (14.9) | 161.7 (13.4) | 100 |
| **Gal β(1-4) GlcNAc (*1-6*)** | **Phi** | **Psi** | **Pop(%)** |
| Cluster 1 | -73.9 (11.5) | -123.5 (16.5) | 98.7 |
| Cluster 2 | -146.2 (8.5) | -142.1 (6.9) | 1.3 |
| **Man α(1-3) Man** | **Phi** | **Psi** | **Pop(%)** |
| Cluster 1 | 71.1 (9.3) | 140.5 (15.6) | 72.9 |
| Cluster 1 | 69.9 (8.8) | 100.63(9.8) | 27.1 |
| **GlcNAc β(1-2) Man (*1-3*)** | **Phi** | **Psi** | **Pop(%)** |
| Cluster 1 | -78.7 (13.8) | 161.1 (12.8) | 90.4 |
| Cluster 2 | -79.1 (7.7) | 110.7 (7.2) | 9.6 |
| **Gal β(1-4) GlcNAc (*1-3*)** | **Phi** | **Psi** | **Pop(%)** |
| Cluster 1 | -72.4 (10.9) | -118.6 (15.2) | 95.0 |
| Cluster 1 | -143.2 (10.4) | -144.4 (5.9) | 2.9 |
| Cluster 1 | -74.2 (9.3) | 67.6 (9.4) | 2.1 |

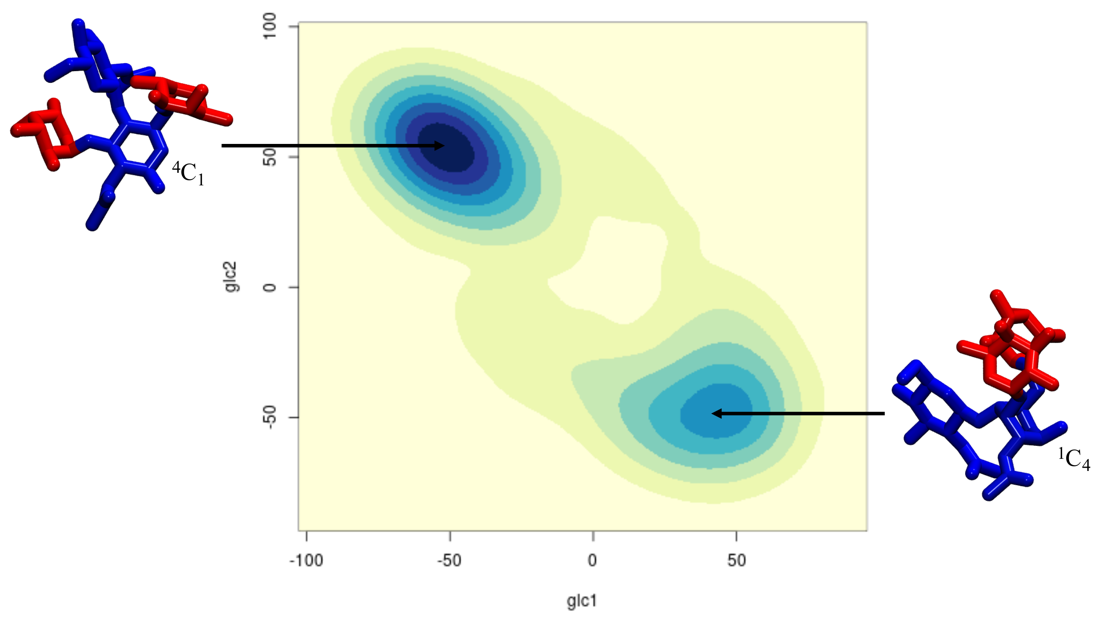

**Figure S.2.** Conformational analysis N-linked GlcNAc pucker along the 1.5 μs cumulative sampling of the α(1-3) and α(1-6) core fucosylated A2G2 (mgmfpf) N-glycan. Representative structures of the fucosylated chitobiose selected from MD sampling of the whole N-glycans are shown on the left- and right-hand side of the heat map where the ring pucker is also indicated. The x and y axis are labelled with the torsion angled measured, namely C1C2C3C4 (glc1) and C2C3C4C5 (glc2), respectively. The monosaccharides colouring follows the SFNG nomenclature. The structure rendering was done with VMD and the graphical statistical analysis with RStudio ([www.rstudio.com](http://www.rstudio.com)).

**Table S.14** Results of the clustering analysis showing the median and standard deviation values (in parenthesis) for the torsion angles (°) measured through a cumulative 2 μs MD sampling of the β(1-2) xylosylated and α(1-3) and α(1-6) core fucosylated mgxmfpf glycan. Note: mg refers to the mammalian terminal β(1-4)-Gal, pf to the plant core α(1-3)-Fuc and mf to the mammalian core α(1-6)-Fuc.

| **Fuc α(1-6) GlcNAc** | **Phi** | **Psi** | **Pop(%)** |
| --- | --- | --- | --- |
| Cluster 1 | -75.9 (11.25) | 175.65 (14.0) | 79.3 |
| Cluster 2 | -142.7 (9.8) | 171.3(6.2) | 16.9 |
| Cluster 3 | -95.48 (4.4) | 71.95(4.8) | 3.8 |
| **Fuc α(1-3) GlcNAc** | **Phi** | **Psi** | **Pop(%)** |
| Cluster 1 | -70.6 (10.3) | 141.4 (8.3) | 88.8 |
| Cluster 2 | -156.0 (8.1) | 89.5 (8.7) | 11.2 |
| **GlcNAc β(1-4) GlcNAc** | **Phi** | **Psi** | **Pop(%)** |
| Cluster 1 | -72.8 (8.9) | -106.6 (11.0) | 96.7 |
| Cluster 2 | -82.1 (7.1) | -154.8 (6.8) | 3.3 |
| **Man β(1-4) GlcNAc** | **Phi** | **Psi** | **Pop(%)** |
| Cluster 1 | -75.5 (11.0) | -123.5 (12.6) | 94.6 |
| Cluster 2 | -62.42 (7.9) | 72.2 (9.5) | 4.4 |
| **Man α(1-6) Man** | **Phi** | **Psi** | **Pop(%)** |
| Cluster 1 | 73.4 (10.5) | 102.8 (10.6) | 55.3 |
| Cluster 2 | 70.4(9.4) | -176.8 (16.7) | 26.7 |
| Cluster 2 | 102.5 (9.3) | 55.9 (7.7) | 17.9 |
| **GlcNAc β(1-2) Man (*1-6*)** | **Phi** | **Psi** | **Pop(%)** |
| Cluster 1 | -91.8 (13.5) | 159.4 (11.1) | 100 |
| **Gal β(1-4) GlcNAc (*1-6*)** | **Phi** | **Psi** | **Pop(%)** |
| Cluster 1 | -74.2 (11.1) | -121.1 (15.3) | 99.2 |
| Cluster 2 | -146.6 (7.5) | -143.6 (4.9) | 0.8 |
| **Man α(1-3) Man** | **Phi** | **Psi** | **Pop(%)** |
| Cluster 1 | 68.7 (9.7) | 115.4 (16.8) | 100 |
| **GlcNAc β(1-2) Man (*1-3*)** | **Phi** | **Psi** | **Pop(%)** |
| Cluster 1 | -79.5 (13.8) | 162.3 (12.4) | 92.4 |
| Cluster 2 | -79.3 (6.9) | 110.3 (6.9) | 7.6 |
| **Gal β(1-4) GlcNAc (*1-3*)** | **Phi** | **Psi** | **Pop(%)** |
| Cluster 1 | -72.6 (11.0) | -119.1 (15.2) | 93.4 |
| Cluster 1 | -143.2 (10.4) | -144.4 (5.9) | 4.9 |
| Cluster 1 | -74.2 (11.6) | 69.7 (12.1) | 1.7 |

**Table S.15** Results of the clustering analysis showing the median and standard deviation values (in parenthesis) for the torsion angles (°) measured through a cumulative 2 μs MD sampling of the β(1-2) xylosylated and α(1-3) fucosylated A2 glycan terminating with LeX on both arms.

| **Fuc α(1-3) GlcNAc** | **Phi** | **Psi** | **Pop(%)** |
| --- | --- | --- | --- |
| Cluster 1 | -70.2 (11.0) | 141.9 (8.4) | 90.8 |
| Cluster 2 | -156.7 (7.6) | 90.2 (8.6) | 9.2 |
| **GlcNAc β(1-4) GlcNAc** | **Phi** | **Psi** | **Pop(%)** |
| Cluster 1 | -72.2 (8.6) | -107.5 (7.7) | 100 |
| **Man β(1-4) GlcNAc** | **Phi** | **Psi** | **Pop(%)** |
| Cluster 1 | -76.4 (15.7) | -124.6 (16.7) | 75.0 |
| Cluster 2 | -68.1 (9.7) | 71.0 (10.1) | 25.0 |
| **Man α(1-6) Man** | **Phi** | **Psi** | **Pop(%)** |
| Cluster 1 | 70.7 (9.2) | -173.6 (13.4) | 89.7 |
| Cluster 2 | 152.4 (12.9) | 145.6 (12.3) | 10.3 |
| **Xyl β(1-2) Man** | **Phi** | **Psi** | **Pop(%)** |
| Cluster 1 | -77.6 (9.6) | 140.1 (18.1) | 100 |
| **GlcNAc β(1-2) Man (*1-6*)** | **Phi** | **Psi** | **Pop(%)** |
| Cluster 1 | -78.2 (14.3) | 162.3 (13.1) | 91.8 |
| Cluster 2 | -76.9 (7.4) | 111.1 (6.8) | 8.2 |
| **Fuc β(1-3) GlcNAc (*1-6*)** | **Phi** | **Psi** | **Pop(%)** |
| Cluster 1 | -69.7 (8.9 | 142.1 (7.0) | 100 |
| **Gal α(1-4) GlcNAc (*1-6*)** | **Phi** | **Psi** | **Pop(%)** |
| Cluster 1 | -67.7 (8.0) | -107.9(7.2) | 100 |
| **Man α(1-3) Man** | **Phi** | **Psi** | **Pop(%)** |
| Cluster 1 | 61.5 (8.5) | 111.0 (15.1) | 100 |
| **GlcNAc β(1-2) Man (*1-3*)** | **Phi** | **Psi** | **Pop(%)** |
| Cluster 1 | -78.8 (13.9) | 162.6 (11.7) | 91.6 |
| Cluster 2 | -79.1 (6.2) | 112.5 (6.2) | 8.3 |
| **Gal β(1-3) GlcNAc (*1-3*)** | **Phi** | **Psi** | **Pop(%)** |
| Cluster 1 | -69.1 (9.45) | 142.2 (8.0) | 100 |
| **Fuc α(1-4) GlcNAc (*1-3*)** | **Phi** | **Psi** | **Pop(%)** |
| Cluster 1 | -67.8 (7.9) | -108.1 (7.4) | 94.8 |
| Cluster 2 | -153.9 (8.8) | -141.5 (7.7) | 5.2 |

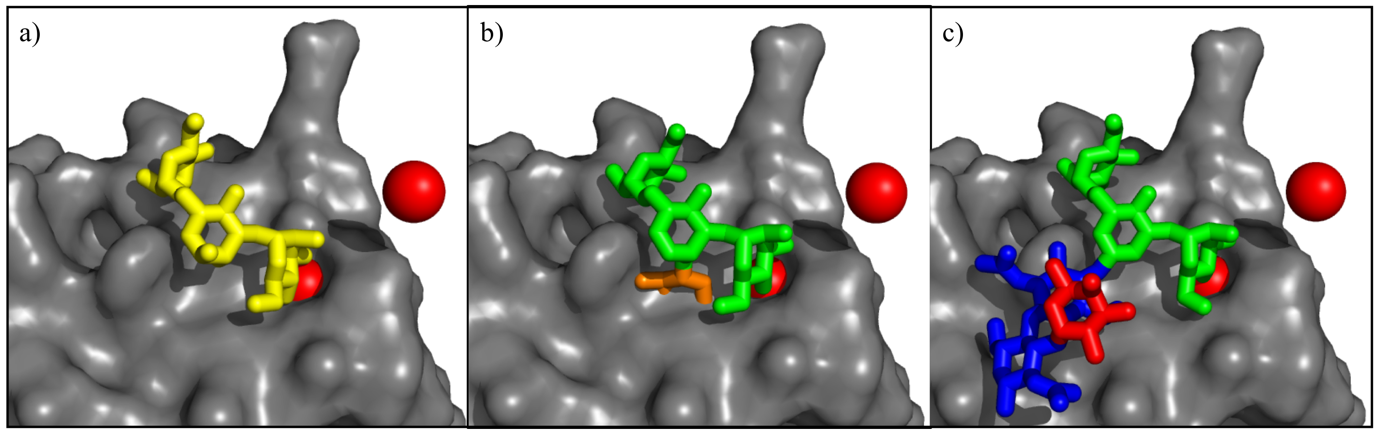

**Figure S.3.** Structural alignment of different plant N-glycoforms from our MD simulations on to the DC-SIGN/GlcNAc2Man3 complex resolved at 2.5 Å resolution (PDBid 1k9i). Panel a) The water accessible surface of the DC-SIGN (chain C) binding site is shown in grey and the Man3 region of the co-crystallized glycan in 1k9i is rendered as yellow sticks. Ca^2+^ ions are shown as red spheres. Panel b) Structural alignment of representative structure from our MD simulation of the ngx plant N-glycan shows that the β(1-6) xylose sterically hinders binding by clashing with the surface of the binding site. Only the Xyl-Man3 glycoblock from the whole N-glycan is represented. Panel c) Structural alignment of representative structure from our MD simulation of the ngf plant N-glycan shows that the α(1-3) fucose does not hinders recognition or binding by DC-SIGN. Only the Man3 and α(1-3)-Fuc chitobiose glycoblocks from the whole N-glycan is represented. The monosaccharides colouring, aside from panel a), follows the SFNG nomenclature. The structure rendering was done with VMD and the graphical statistical analysis with RStudio ([www.rstudio.com](http://www.rstudio.com)).

1. These authors have contributed equally to the work
